# Measles-based Zika vaccine induces long-term immunity and requires NS1 antibodies to protect the female reproductive tract in the hCD46 IFNα/β receptor knockout mice

**DOI:** 10.1101/2020.09.17.301622

**Authors:** Drishya Kurup, Christoph Wirblich, Matthias J. Schnell

## Abstract

Zika virus (ZIKV) can cause devastating effects in the unborn fetus of pregnant women. To develop a candidate vaccine that can protect human fetuses, we generated a panel of live measles vaccine (MV) vectors expressing ZIKV-E and -NS1. Our MV-based ZIKV-E vaccine, MV-E2, protected mice from the non-lethal Zika Asian strain (PRVABC59) and the lethal African strain (MR766) challenge. Despite 100% survival of the MV-E2 mice, however, complete viral clearance was not achieved in the brain and reproductive tract of the lethally challenged mice. We then tested a combination of two MV-based vaccines, the MV-E2 and a vaccine expressing NS1 (MV-NS1[2]), and we observed durable plasma cell responses, complete clearance of ZIKV from the female reproductive tract, and complete fetal protection in the lethal African challenge model. Our findings suggest that NS1 antibodies are required to enhance the protection achieved by ZIKV-E antibodies in the female reproductive tract.

## Introduction

Zika virus (ZIKV), an emerging mosquito-borne pathogen, in most healthy adults causes only mild infection, but in rare cases, causes Guillain-Barré syndrome (GBS) in adults and congenital Zika syndrome (CZS) in infants born to ZIKV-infected mothers (*1*). ZIKV is primarily transmitted by mosquitoes of the *Aedes* genus, but can also be transmitted through congenital, perinatal, blood transfusion, and sexual routes (*2*). Before the 2007 ZIKV outbreak in the Yap Islands, only 14 human cases were reported worldwide (WHO, Accessed November 12, 2019). Since then, outbreaks have occurred in the Pacific islands, South and Central America, and the Caribbean (WHO, November 12, 2019). These unprecedented outbreaks led to a sudden increase in human cases with large numbers of symptomatic infections characterized by fever, conjunctivitis, rash, headache, myalgia, and arthralgia (*3*). Also, retrospective studies of the epidemic showed a strong correlation of ZIKV disease with microcephaly and/or other congenital disabilities in infants and GBS in adults (*4*). Based on these case studies, WHO declared ZIKV as a Public Health Emergency of International Concern on Feb 1, 2016 (WHO, Accessed November 12, 2019).

ZIKV belongs to the *Flavivirus* genus of the *Flaviviridae* family. ZIKV contains a single-stranded positive-sense RNA genome containing a 5’ untranslated region (UTR), a single open reading frame (ORF) encoding a polyprotein, and a 3’ UTR. The ORF encodes three structural proteins (capsid, C; pre-membrane, prM; and envelope, E) and seven nonstructural proteins (NS1, NS2A, NS2B, NS3, NS4A, NS4B, NS5) (*5*). The prM protein associates with E to form heterodimers and is important for the proper folding of E (*6*). Among the structural proteins, the E protein is the major virion surface protein, but the M protein is also displayed on the surface of the viral particle. C protein is a major internal protein that is surrounded by the host-derived spherical lipid bilayer membrane (*7*). The glycosylated NS1 protein forms a homodimer and separates into three distinct populations: a large portion localizes to the site of viral RNA synthesis and is critical for replication; a second minor portion traffics to the plasma membrane (mNS1, membrane) where it forms a hydrophobic “spike”, which may contribute to its cellular membrane association; and a third is secreted into the extracellular space as a hexamer (sNS1, secreted) (*8*). The sNS1 is secreted in the serum of infected individuals in high concentrations and is used as a diagnostic biomarker (*9*). Currently, there are two distinct strains of ZIKV, the Asian and African, but only one serotype (*10*). The Asian strain associated with the recent outbreaks evolved from the first isolated 1947 African strain after sporadic cases of ZIKV in Africa and Asia (WHO, November 12, 2019).

ZIKV infection in healthy adults generates virus neutralizing antibodies (VNAs) directed towards the E protein and antibodies directed towards the NS1 protein. The presence of ZIKV-E and NS1 antibodies are suggestive of protective immunity in humans (*11, 12*). While the protective E protein antibodies are neutralizing and target the virions, the NS1 antibodies are non-neutralizing and target the infected cells (*13*). Several candidate vaccines have utilized the ZIKV ME, prME, and/or NS1 as the immunogen of choice in DNA, RNA, viral vectors, live attenuated vaccine (LAV), inactivated virus, and subunit vaccine platforms (*13-16*). Some of these vaccines have proceeded to Phase I/II clinical trials that demonstrated safety and immunogenicity (*17-23*). When these vaccine platforms were tested in non-pregnant mice and monkeys, they achieved systemic viral clearance, however, the vaccines tested in pregnant mouse and monkey models achieved incomplete fetal protection (*24, 25*). Recently, the RhAd52- and Ad26-based ZIKV vaccine tested in pregnant *IFNαβR*^*-/-*^ mice were more successful: just marginal levels of ZIKV RNA were detected in the placenta and fetal brains (*26*).

Each of these vaccine platforms has its advantages and shortcomings. The DNA, RNA, VLP, and subunit vaccine are likely safe platforms that require multiple doses, but the longevity of the vaccine-induced immune responses in humans is unknown. Viral vectors, like modified vaccinia virus Ankara (MVA) and vesicular stomatitis virus (VSV), induce durable responses but have safety implications when administered to children < 1 year of age, pregnant women, and the immunocompromised (*27, 28*). While protective parameters are yet to be established for ZIKV, the development of a certain threshold of neutralizing E protein antibodies is considered protective for other flaviviruses, such as yellow fever (YF), tick-borne encephalitis virus (TBEV), and Japanese encephalitis virus (JEV). But ZIKV, unlike other flaviviruses, can cause devastating effects in pregnant women, resulting in prolonged viremia leading to CZS in their infants (*29*). Hence a ZIKV vaccine must be able to prevent viremia in pregnant women and their fetuses.

In an attempt to safely and effectively overcome the persistence of viremia in pregnant women and the impact on their fetuses, we developed a ZIKV vaccine based on the measles vaccine vector. The measles virus (MV) vaccine that has more than 50 years of historical data confirming its safety and long-term efficacy, by induction of durable neutralizing antibody and T cell-mediated immunity (*30, 31*). Children < 1 year of age, pregnant women, and postpartum women can be MV vaccinated (*30, 32, 33*). The success of the MV vaccine has led to its development as a vaccine vector for DENV, WNV, HIV, Middle East respiratory syndrome (MERS), and malaria antigen (*30*).

Here, we used the MV vaccine vector (Edmonston B strain) to generate a candidate ZIKV vaccine that can be included in the childhood vaccination regime. We inserted the codon optimized (co) ZIKV prME from the Asian PRVABC59 strain into the transcription cassette before N (position 0) and another between N and P (position 2), generating the MV-E(0) and the MV-E(2) vaccine, respectively. Both vaccines completely cleared the virus in mice challenged with the non-lethal ZIKV PRVABC59 (Asian) 2015 strain and dramatically reduced ZIKV RNA copies when challenged with the lethal highly neurotropic mouse-adapted ZIKV MR766 (African) strain. Second-generation vaccine constructs containing the ZIKV NS1 protein-MV-NS1(2) were tested singly or in combination with MV-E(2) or by inserting the co ZIKV prMENS1 between H and L protein (position 6), to allow for enhanced protection using the lethal mouse-adapted ZIKV African MR766 strain. Our results showed that NS1 antibodies alone did not protect as the MV-NS1(2) mice succumbed to the lethal Zika African strain challenge. Interestingly, the combination of MV-NS1(2) and MV-E2 virus provided better protection than MV-E2 alone, in terms of neutralizing titers and clearing ZIKV RNA in the female reproductive tract. In addition, fetuses born to pregnant mice vaccinated with the combination of MV-NS1(2) and MV-E2 vaccines were completely protected from the lethal ZIKV African strain challenge. Lastly, the combination vaccine also induced ZIKV-E-, ZIKV-NS1-, and MV-H-specific long-lived and short-lived plasma cell responses. These findings suggest that further development of this vaccine could lead to an effective pre-exposure Zika vaccine for children.

## Results

### Design, recovery, and characterization of first-generation MV-ZIKV vaccines

Recombinant measles viruses (rMV) have been used as vaccine vectors for different infectious diseases and can serve as an excellent vaccine platform to generate an early childhood vaccine for ZIKV. A full-length measles virus cDNA clone (MV-ATS-0) that allows the insertion of foreign genes upstream of the nucleoprotein gene served as the backbone of our vaccine constructs. We modified the cDNA clone by the addition of a hammerhead ribozyme before the leader region. We generated two additional vectors, MV-ATS-2 and MV-ATS-6, that allow the insertion of foreign genes between the N and P genes and the H and L genes, respectively. We chose to evaluate multiple vectors because transcription of the measles virus genome occurs sequentially, which results in a transcription gradient as the polymerase proceeds from one gene to the next. The point of insertion also affects the replication and spread of the recombinant viruses. We first inserted the codon-optimized (co) gene of the ZIKV precursor (pr), membrane (M), and envelope (E) protein (strain PRVABC59 Asian 2015, Supplemental Fig. 1) upstream (ATS-0) and downstream (ATS-2) of the nucleoprotein gene (Figure 1A). The recombinant viruses rMV-E0 and rMV-E2 were recovered as described in Materials and Methods and amplified on VERO cells. Control viruses that express a green fluorescent protein (GFP) at ATS-0 and a GFP-nanoluciferase fusion gene at ATS-2 were generated in a similar way (Figure 1A). Characterization of the rMV-ZIKV viruses was performed by an immunofluorescence assay to examine ZIKV-E and MV nucleoprotein (N) expression (Fig. 1B). Vero cells infected at a MOI of 0.1 for three days were permeabilized and stained with antibodies directed against ZIKV-E and MV-N. Co-expression of MV-N and ZIKV-E was detected for the MV-E0, and MV-E2 viruses with only ZIKV-E expressed in the ZIKV PRVABC59-infected cells. MV-N but not ZIKV-E was detected in the empty vector rMV infected cells.

**Fig. 1.**
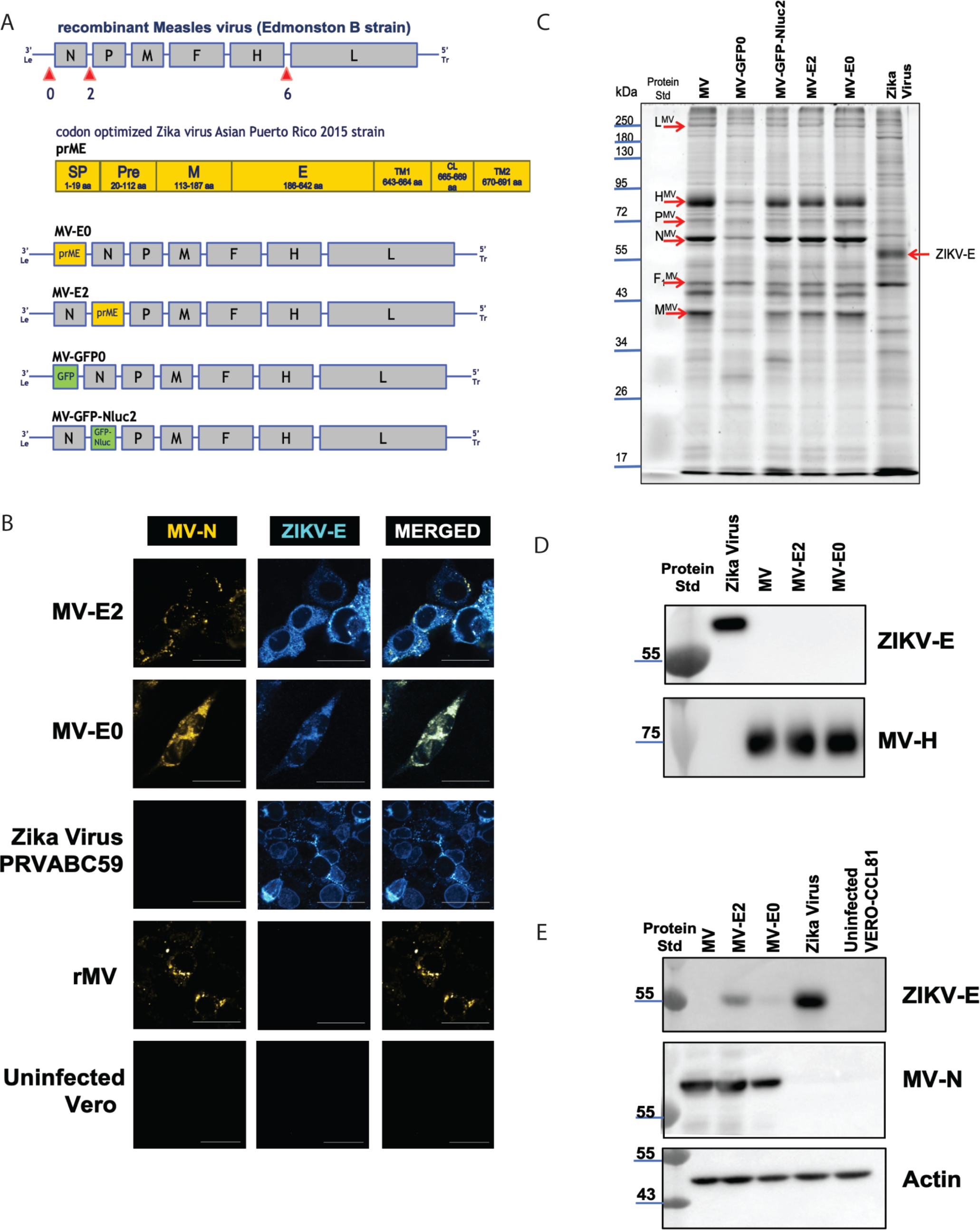
Generation and characterization of first-generation MV-based candidate ZIKV vaccines. **(A)** MV-ZIKV vaccine constructs and controls. **(B)** Immunofluorescence staining of Vero cells was infected at MOI-0.1 for 72 hours with the recovered MV-ZIKV candidate vaccines and control viruses. The permeabilized cells were stained for MV using an α-MV nucleoprotein mouse monoclonal and for the ZIKV-E using a mouse monoclonal antibody. The MV-N antibody is conjugated with a Dylight 488 (green color changed to **Yellow** for visualization) fluorophore, and ZIKV-E is stained with a secondary goat α-mouse Cy-3 (red color changed to **Cyan** for visualization) antibody. Confocal images were taken using NIKON-A1R, 60X magnification, 3X zoom. The scale bar measures 30 μm. **(C)** Sucrose-purified virions were analyzed on SDS-PAGE (10%) stained with SYPRO Ruby. Zika virus PRVABC59 strain was loaded as the control. **(D)** Western blot analysis of sucrose-purified virions probed for ZIKV-E and MV-H. Zika virus PRVABC59 strain was loaded as the control. **(E)** Western blot analysis of cell lysates of VERO cells infected with MV-ZIKV candidate vaccine at MOI-5 for 60 hours. Zika virus PRVABC59 infected cell lysates were loaded as the control. The blot was probed for ZIKV-E and MV-N.

Next, we purified the rMV-ZIKV viruses over a 20% sucrose-cushion and analyzed by SDS-PAGE (10%). The Sypro-ruby stained gel showed the presence of all six MV structural proteins in the MV-E0 and MV-E2 viruses migrating at a similar size compared to the empty vectors rMV, MV-GFP0, and NV-GFP-Nluc2 viruses (Fig. 1C). Western blot analysis of a similar SDS-PAGE of purified rMV-ZIKV vaccines probed for ZIKV-E showed the absence of ZIKV-E in the virion (Fig. 1D), indicating that rMV-ZIKV vaccines do not incorporate the ZIKV E protein, thereby retaining the tropism of the MV vector. Cell lysates obtained from Vero cells infected by rMV-ZIKV vaccines at a MOI of 5 for 60 hours were analyzed on a western blot. The western blot probed for ZIKV-E & MV-N confirmed their expression (Fig. 1E) in both the MV-E0 and MV-E2 viruses, with lower levels of ZIKV-E, expressed by the MV-E0 virus. Similar levels of the actin control were detected in the western blot of the cell lysates (Fig. 1E).

### Efficacy of MV-ZIKV vaccines using non-lethal ZIKV Asian PRVABC59 strain

The rMV vaccine strain requires both the presence of the human CD46 receptor (hCD46) for its replication (*34*) and the lack of the interferon αβ receptor (*IFNαβR*^*-/-*^) for its systemic replication in mice (*35*). Therefore, rMV vaccines historically have been tested in *hCD46 IFNαβR*^-/-^ transgenic mice. To initially test the immunogenicity of our new vaccines, two groups consisting of five female *hCD46 IFNαβR* ^*-/-*^ mice (10-to 12-weeks old) each were immunized with either MV-E2 or MV-E0 on day 0 and boosted on day 28 with 10^5^ TCID_50_ intraperitoneally (i.p.); two additional groups were mock immunized with PBS at the same time points (Fig. S2A). The mice were bled on days 0, 28, 35, and 63 and tested for the presence of ZIKV-E and MV-H IgG antibody by ELISA.

After the boost, the MV-E2 vaccinated mice showed significantly higher titers against ZIKV-E (p = 0.002) in comparison to the MV-E0 vaccinated mice (Fig. S2B). By day 63, the MV-E0 and MV-E2 vaccinated mice had similar ZIKV-E EC_50_ IgG titers (Fig. S1B). The MV-H responses were similar for the MV-E0, and the MV-E2 vaccinated animals for all time points (Fig. S2C). Since the vaccines showed strong immunogenicity, the mice were challenged with the 10^6^ FFU of non-lethal ZIKV Asian PRVABC59 strain subcutaneously (s.c.) on day 63 to mimic the natural infection route; they were then humanely euthanized at day 77 (14 days post-challenge). One PBS group was mock challenged with PBS and served as a control.

The MV-E2 and MV-E0 vaccines provide robust protection with undetectable ZIKV RNA in the blood of vaccinated animals, while the control PBS group had significantly high RNA copies ∼10^4^-10^7^ in the blood on day 7 and 14 post challenge (Fig. S2E). Similar results were observed in the brain (Fig. S2F) and the reproductive tract (Fig. S2G). The high ZIKV neutralizing titers seen before the challenge (Fig. S2D) were maintained at the necropsy time point (Fig. S2H) for both the MV-E2 and the MV-E0 groups.

### Efficacy of MV-ZIKV vaccines using lethal mouse-adapted ZIKV African MR766 strain

To test whether the MV-ZIKV vaccines could be efficacious in a lethal mouse challenge model in both males and females, eight groups of five female (F) or male (M) *hCD46 IFNαβR* ^-/-^ mice (7-to 8-weeks old) each were immunized with 10^5^ TCID_50_ i.p. of either MV-E2(F), MV-E2(M), MV-E0(F), MV-E0(M), MV-GFP0(M), MV-GFP-Nluc2(M), rMV(F), or rMV(M) vaccine, on day 0 and boosted on day 21 (Fig. 2A). The mice were bled on days 0, 14, 28, 56, and 104 and tested for the presence of ZIKV-E and MV-H IgG antibody titers.

**Fig. 2.**
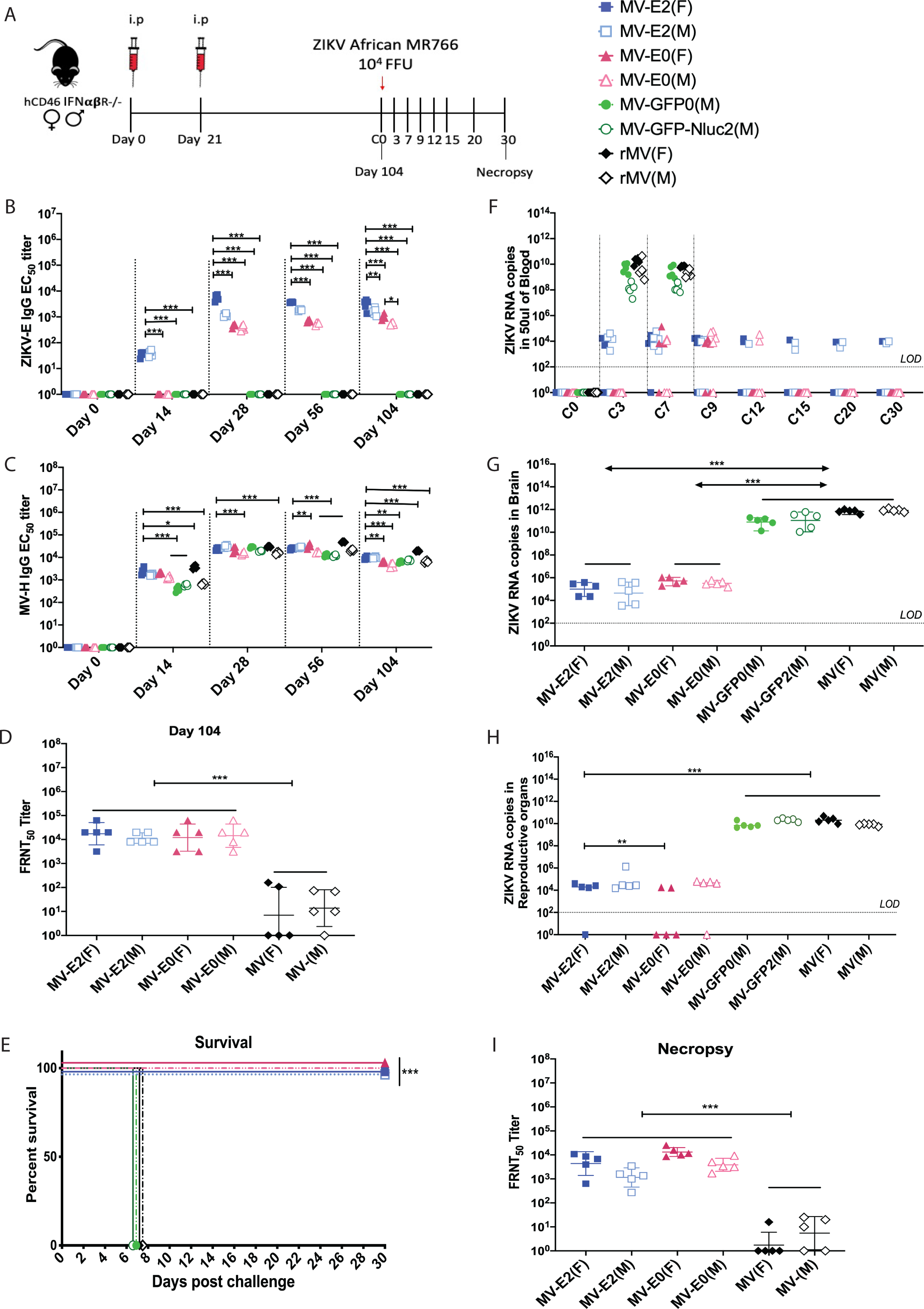
Immunogenicity and efficacy testing of first-generation candidate MV-ZIKV vaccines using a lethal ZIKV African MR766 challenge strain. **(A)** Timeline of vaccination, challenge, and viral load determinations. **(B-C)** anti-ZIKV-E-specific (B), and anti-MV-H specific (C) ELISA EC_50_ titers of the vaccinated animals are plotted on a graph for all animals at different time points. The Mean ± SD is depicted per group. **(D & I)** ZIKV neutralization with PRVABC59 Asian strain. FRNT was performed on day 104 (D) and necropsy (I) sera from vaccinated animals and controls. The 50% neutralizing titer (FRNT_50_) is plotted on the graph. The Mean ± SD is depicted per group. **(E)** Kaplan-Meier survival curve analysis of vaccinated and control animals post challenge. **(F-H)** ZIKV RNA copies by quantitative polymerase chain reaction (qPCR) in the blood (F), brain (G), and reproductive tract (H). The Mean ± SD is depicted per group. Statistics for Fig. 2B-D & F-I were done using the two-way ANOVA with posthoc Tukey HSD test and performe on log-transformed data for each time point. Fig. 2E, Survival curves were analyzed using the log-rank test with a Bonferroni correction. Only significant differences are depicted. P-value of 0.12(ns), 0.033(*), 0.002(**), <0.001(***) are depicted accordingly. A horizontal line () is used to include all groups below it.

The ZIKV-E immune responses developed as early as day 14 in the MV-E2 group, while for the MV-E0 group, detectable responses were seen only after the boost on day 28. The MV-E2 vaccinated mice developed significantly higher ZIKV-E-specific antibody titers than the MV-E0 vaccinated animals at all time points. This difference was only observed in younger mice. Significantly higher ZIKV-E IgG antibody titers were observed in the females than in the males in both MV-E2 and MV-E0 vaccinated groups (Fig. 2B). The MV-H IgG responses were the highest in the rMV(F) at all time points, with higher MV-H antibody titers seen in the females than the males of the empty MV vector group. The MV-E2 group developed significantly higher MV-H IgG antibody titers than the MV-E0, MV-GFP0, and the MV-GFP-Nluc2, indicating that the vector immunity was unaffected by the addition of the ZIKV-E into the genome (Fig. 2C).

The vaccinated animals were challenged on day 104 with a lethal dose of 10^4^ FFU s.c. of the lethal ZIKV African MR766 strain. All of the MV-E2 and MV-E0 vaccinated male and female mice survived the challenge, showing no signs of ZIKV disease, while the MV-GFP0(M), MV-GFP-Nluc2(M), and rMV(F&M) controls succumbed to ZIKV disease by day 7 and were humanely euthanized (Fig. 2E). Significantly lower ZIKV RNA copies were observed in the blood of the MV-E2 and MV-E0 groups than of the control groups (MV-GFP0, MV-GFP-Nluc2, and rMV) at all time points, with complete viral clearance from the blood seen in the MV-E0 vaccinated animals by day 15 (Fig. 2F). Similar to the viral load in the blood, significantly lower ZIKV RNA copies were observed in the brain (Fig. 2G) and the reproductive tract (Fig. 2H) of the MV-E2 and the MV-E0 vaccinated animals than of the control groups. The MV-E2 and MV-E0 vaccinated animals developed similar ZIKV neutralizing titers at day 104 and necropsy (Fig. 2D and I).

### MV-ZIKV vaccines induce long-term immunity and protection

We next assessed the longevity of the immune responses induced by MV-E2 and MV-E0 vaccines. Four groups of five female *hCD46 IFNαβR* ^*-/-*^ mice (9-to 12-weeks old) were vaccinated on day 0 and boosted on day 28 with 10^5^ TCID_50_ i.p. of MV-E2, MV-E0, rMV, or PBS (Fig. S3A). The MV-E2 and MV-E0 groups had similar ZIKV-E IgG antibody titers by day 110 (Fig. S3B). Similar to the previous experiment, MV-H IgG antibody titers were similar in the MV-E2-, MV-E0-, and the MV-vector vaccinated animals (Fig. S3C). Of note, MV-E2 and MV-E0 vaccinated animals survived 10^4^ FFU of lethal ZIKV African MR766 strain challenge (Fig. S3E) with viral clearance in the blood at necropsy (Fig. S3F) and significantly lower ZIKV RNA copies in the brain (Fig. S3G) and the reproductive tract compared to the control animals (Fig. S3H). Viral clearance correlated with high ZIKV neutralizing titers on day 144 (Fig. S3D) and at necropsy (Fig. S3I).

### MV-E2 vaccine is efficacious when administered intramuscularly

To learn whether the route of immunization is important and whether prior MV immunity affected the efficacy of the MV-E2 and MV-E0 vaccines, we performed an additional mouse challenge study. Four groups of five male or female *hCD46 IFNαβR* ^-/-^ mice (9-to 12-weeks old) were pre-vaccinated (Prevac) on day –35 with 10^5^ TCID_50_ intramuscularly (i.m.) of rMV. Then, on day 0, they were vaccinated i.m. with 10^5^ TCID_50_ of either MV-E2 or MV-E0 vaccines and boosted on day 21 (Fig. S4A). In addition, on day 0, seven groups of five female or male mice were vaccinated i.m. with MV-E2, MV-E0, MV, or PBS. The ZIKV-E IgG titers followed a similar trend to the i.p. MV-E2 and MV-E0 vaccinated animals, but lower titers were seen in the i.m. vaccinated animals (Fig. S4B). For the Prevac groups, MV-E2 vaccinated animals elicited significantly lower ZIKV-E IgG titers while the MV-E0 vaccinated animals did not seroconvert throughout the study (Fig. S4B). The MV-H IgG titers were boosted on day 0 and 21 in the Prevac groups, which indicated successful vaccination. The Prevac groups developed similar MV-H antibody responses by day 56 to the MV-E2, MV-E0, and rMV groups. All the animals were challenged on day 63 with 10^4^ FFU of lethal ZIKV African MR766 strain. All of the MV-E2 vaccinated animals survived the challenge. In contrast, the animal with the lowest ZIKV neutralizing titer (Fig. S4D) in the MV-E0 group succumbed to challenge, confirming that a certain threshold of neutralizing titer determines protection (Fig. S4E). All of the mice in the Prevac groups succumbed to ZIKV disease on day 9, while controls succumbed on day 7. The Prevac groups and the control groups showed significantly higher ZIKV RNA in the blood, brain, and the reproductive tract than the MV-E0 and MV-E2 vaccinated animals (Fig. S4F-H). The findings of this study confirm that MV-E2 vaccine is efficacious irrespective of the route of immunization (Fig. 3, S2-S4), and that prior MV immunity affects the efficacy of the MV-E2 and MV-E0 vaccines.

**Fig. 3.**
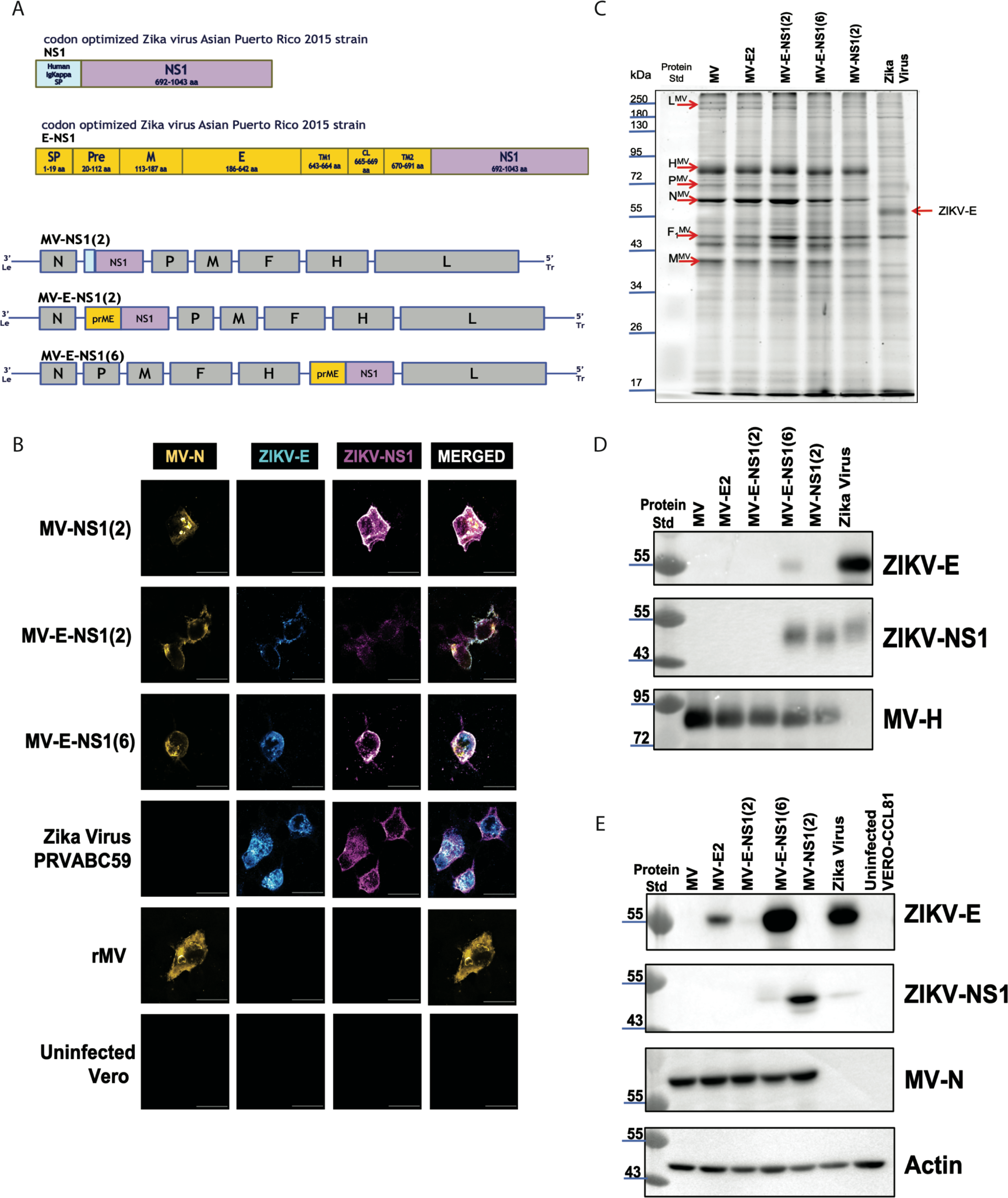
Generation and characterization of second-generation MV-based candidate ZIKV vaccines. **(A)** Second-generation vaccine constructs **(B)** Immunofluorescence staining of Vero cells was infected at MOI-0.1 for 72 hours with the recovered modified MV-ZIKV candidate vaccines and control viruses. The permeabilized cells were stained for MV using an α-MV N protein mouse monoclonal conjugated with dylight488 (green color changed to **Yellow** for visualization), for the ZIKV-E using a mouse monoclonal antibody, and for ZIKV NS1 using human monoclonal antibody EB9. ZIKV-E is stained with a secondary α-mouse AF568 (red color changed to **Cyan** for visualization) antibody, and ZIKV-NS1 is stained with a secondary goat α-human AF-647 (far-red color changed to **Magenta** for visualization) antibody. Confocal images were taken using NIKON-A1R, 60X magnification, 2.5X zoom. The scale bar measures 30 μm. **(C)** Sucrose-purified virions were analyzed on SDS-PAGE (10%) stained with SYPRO Ruby. Zika virus PRVABC59 was loaded with the control virus **(D)** Western blot analysis of sucrose purified virions probed for ZIKV-E and MV-H. Zika virus PRVABC59 was loaded as the control virus. **(E)** Western blot analysis of cell lysates of Vero cells infected with the MV-ZIKV candidate vaccine at MOI-5. Zika virus PRVABC59 infected cell lysates were loaded as the control. The blot was probed for ZIKV-E, ZIKV-NS1, and MV-N.

### Rationale, design, and characterization of second-generation MV-ZIKV vaccines

While the MV-E2 vaccine was efficacious when administered i.p. or i.m., it did not completely protect the brain and the reproductive tract of vaccinated animals when challenged with the mouse-adapted ZIKV African MR766 strain. ZIKV NS1 is expressed on the surface of infected cells, and previous research by others has shown that antibodies directed towards NS1 are protective via antibody-dependent cellular cytotoxicity (ADCC) or antibody-dependent cellular phagocytosis (ADCP) (*13*). We, therefore, generated MV-NS1(2) that expressed ZIKV NS1 from ATS-2 and swapped the ZIKV signal peptide (SP) with that of the human Ig Kappa signal peptide to allow for better secretion (Supplemental Fig. S5) (*13*). Additionally, we generated constructs that expressed the prME-NS1 (Supplemental Fig. S6) from ATS-2 and ATS-6, naming them MV-E-NS1(2) and MV-E-NS1(6), respectively, to test whether the NS1 antibodies enhanced the protection elicited by ZIKV-E antibodies (Fig. 3A). We recovered these second-generation MV-ZIKV vaccines using the standard methods described in the Materials and Methods section.

The recovered second*-*generation MV-ZIKV viruses were assessed for their expression of ZIKV-E, ZIKV-NS1, and MV-N by immunofluorescence assay (Fig. 3B). The MV-NS1(2) co-express MV-N and ZIKV-NS1, while MV-E-NS1(2) and MV-E-NS1(6) express MV-N, ZIKV-E, and ZIKV-NS1 proteins. The control ZIKV PRVABC59-infected cells stained for ZIKV-E and ZIKV-NS1 while the empty vector rMV (Edmonston B strain) only stained for MV-N.

To investigate the influence of the foreign gene on virus production, we assessed multi-step virus growth kinetics of MV-ZIKV vaccines (Supplemental Fig. S7). Released and cell-associated viruses were harvested at all time points. The MV-E2 virus showed peak titers similar to the empty MV vector, while the MV-NS1 yielded slightly lower titers. In contrast, the MV-E0 yielded very low titers. For the single antigen (ZIKV-E or NS1) viruses, peak titers were seen at 120 hpi, while for the dual antigen vaccines, peak titers were seen at 96 hpi. The MV-E-NS1(6) grew to higher titers than MV-E-NS1(2).

Next, the rMV-ZIKV viruses were purified over a 20% sucrose-cushion and analyzed by SDS-PAGE (10%). The Sypro-ruby stained gel showed the presence of all six MV structural proteins in the MV-NS1(2), MV-E-NS1(2), and MV-E-NS1(6) viruses, similar to empty vector rMV and MV-E2 viruses (Fig. 3C). Western blot analysis of the similar SDS-PAGE of purified rMV-ZIKV vaccines probed for ZIKV-E showed an absence of ZIKV-E in all constructs except MV-E-NS1(6) (Fig. 3D) in the virion. The ZIKV-E and -NS1 may sometimes co-purify with the virions, as seen by the slight ZIKV-E band in MV-E-NS1(6), as well as NS1 bands seen in the MV-E-NS1(6), MV-NS1(2), and Zika virus virions (Fig. 3D). The western blot of sucrose purified virions showed the presence of MV-H in all MV’s except for the control Zika virus virions.

Cell lysates obtained from Vero cells infected by rMV-ZIKV vaccines at a MOI of 5 for 60 hours were probed for ZIKV-E, ZIKV-NS1, and MV-N in a western blot (Fig. 3E). The western blot confirmed the expression of ZIKV-E in MV-E-NS1(6) and MV-E2, with greater expression seen for the MV-E-NS1(6). A faint ZIKV-E band is seen in the MV-E-NS1(2) cell lysates, which correlates with its slow replication (Fig. 3E, Supplemental Fig. S7). The ZIKV-NS1 expression is the highest in the MV-NS1(2) cell lysates, with low levels seen in the MV-E-NS1(6) and the control Zika virus infected cell lysates. The low-levels of ZIKV-NS1 seen may be due to the efficient secretion of the NS1 out of the cells. No ZIKV-NS1 was seen in the MV-E-NS1(2) cell lysates. A similar MV-N expression was seen in all the recombinant MV cell lysates (Fig. 3E). Similar levels of the actin control were detected in the western blot of the cell lysates (Fig. 3E).

Next we wanted to characterize the ZIKV-E and ZIKV-NS1 protein secreted by our MV-ZIKV vaccines. The purified SVPs resuspended in non-reducing buffer were probed for ZIKV-E and ZIKV-NS1 in a western blot (Supplemental Fig.S8). The ZIKV-E probed blot showed the presence of a monomeric and a dimeric band similar to the Zika virus made SVP, in the MV-E2 and the MV-E-NS1(6) lane, while no band is seen in the controls– empty MV, MV-NS1(2) lanes. In addition, a faint-strong band above 250 kDa was also seen in MV-E2, MV-E-NS1(6), and Zika virus lanes. This suggests that the 1^st^ and 2^nd^ generation MV-ZIKV vaccine generates SVPs similar to the Zika virus. The blot probed for NS1 yielded a smear ranging from 55kDa to ∼70kDa above the size of the monomeric NS1 (50kDa), indicating that the NS1 made by the MV-ZIKV vaccines was in a combination of monomeric and dimeric (membrane bound) forms. SVPs for the MV-E0 virus could not be purified and were therefore not included in this blot.

### Efficacy of second-generation MV-ZIKV vaccines using lethal mouse-adapted ZIKV African MR766 strain

Six groups of five female *hCD46 IFNαβR* ^-/-^ mice (10-to 12-weeks old) each were immunized with 10^5^ TCID_50_ i.p. on day 0 and boosted on day 21 with either MV-NS1(2), combination vaccine group—MV-E2 & MV-NS1(2), MV-E-NS1(6), MV-E2, and two PBS groups (Fig. 4A). The combination vaccine group received 10^5^ TCID_50_ each of MV-E2 & MV-NS1(2) vaccine (2 × 10^5^ TCID_50_ total) (Fig. 4A). The MV-E-NS1(2) was not included in this study because a pilot study with it failed to achieve seroconversion in mice, apparently due to its slow replication (Supplemental Fig. S7). For all time points, the combination vaccine group elicited similar ZIKV-E IgG antibody titers to the MV-E2 vaccine group, with only modestly higher ZIKV-E responses than the MV-E-NS1(6) vaccinated animals (p = 0.05) (Fig. 4B). The MV-NS1(2) and the MV-E-NS1(6) elicited similar ZIKV-NS1 antibody titers, while the combination vaccine group had lower ZIKV-NS1 responses (Fig. 4C). The combination vaccine group had similar MV-H responses to the MV-E2 group, but significantly higher titers than the MV-NS1(2) and the MV-E-NS1(6) groups (Fig. 4D).

**Fig. 4.**
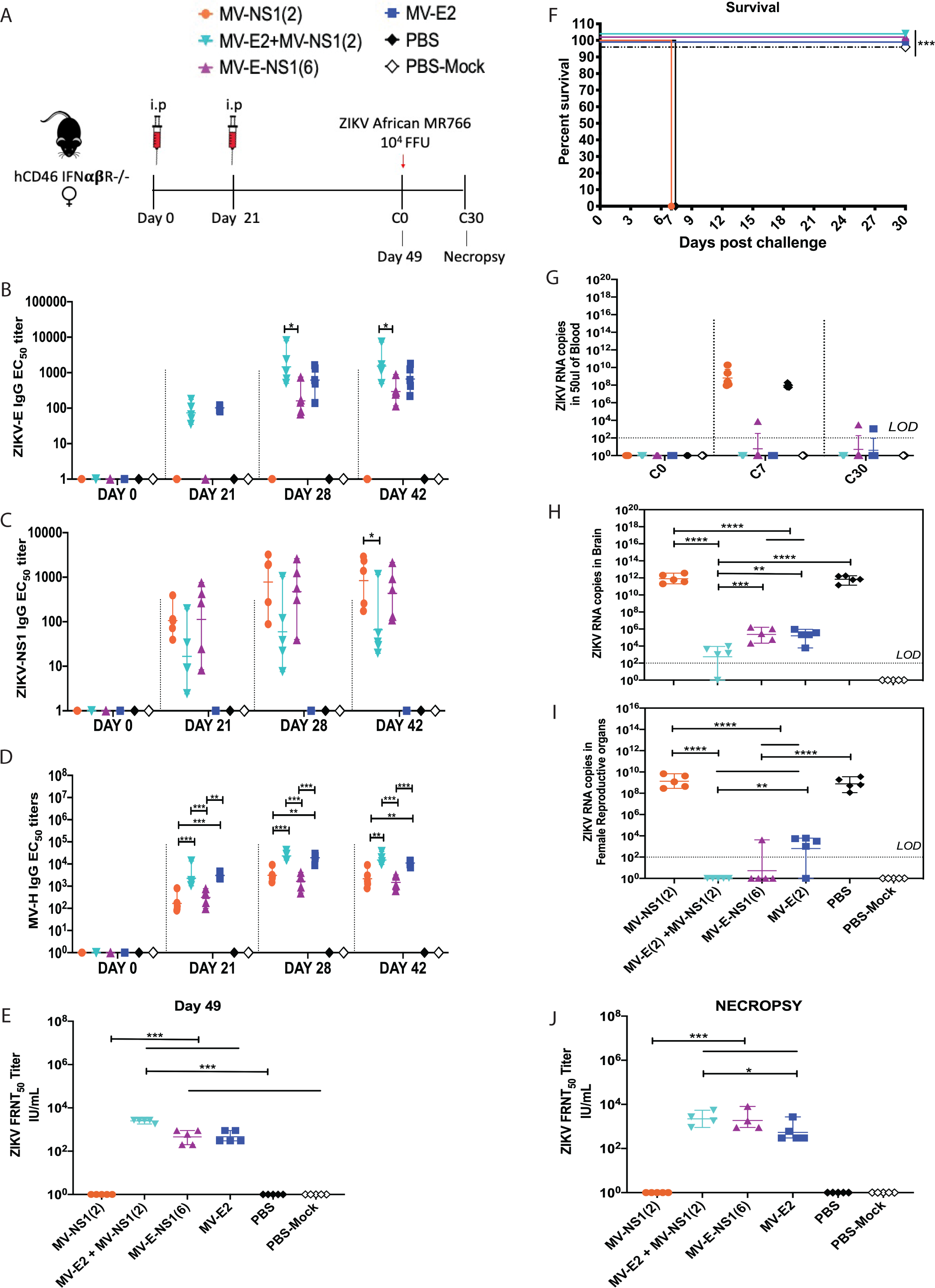
Immunogenicity and efficacy testing of second-generation candidate MV-ZIKV vaccines using a lethal ZIKV African MR766 challenge strain. **(A)** Timeline of vaccination, challenge, and viral load determinations. **(B-D)** anti-ZIKV-E (B), anti-ZIKV-NS1 (C), and anti-MV-H (D) ELISA EC_50_ titers of the vaccinated animals are plotted on a graph for all animals at different time points. The Mean ± SD is depicted per group. **(E & J)** ZIKV neutralization with PRVABC59 Asian strain. FRNT was performed on day 49 (E) and necropsy (J) sera from vaccinated animals and controls. The 50% neutralizing titer (FRNT_50_) is plotted on the graph in terms of IU/mL. The Mean ± SD is depicted per group. **(F)** Kaplan-Meier survival curve analysis of vaccinated and control animals post challenge. **(G-I)** ZIKV RNA copies by qPCR in the blood (G), brain (H), and reproductive tract (I). The Mean ± SD is depicted per group. Statistics for Fig. 4B-E, G-J were done using the one-way ANOVA with posthoc Tukey HSD test and performed on log-transformed data for each time point. Fig. 4E, Survival curves were analyzed using the log-rank test with a Bonferroni correction. Only significant differences are depicted. P-value of 0.12(ns), 0.033(*), 0.002(**), <0.001(***) are depicted accordingly. LOD stands for limit of detection. A horizontal line () is used to include all groups below it.

Based on the collected immunogenicity data, we decided to challenge the mice on day 49 with 10^4^ FFU of ZIKV African MR766 strain s.c., with the exception of one PBS group that was mock challenged with PBS. As expected, the combination vaccine group MV-E2 & MV-NS1, the MV-E-NS1(6) group, and the MV-E2 group survived the challenge, showing no signs of ZIKV disease (Fig. 4F). No detectable ZIKV RNA was seen in the blood of the combination vaccine group, while one animal in the MV-E2 and the MV-E-NS1(6) groups showed the presence of ZIKV that was cleared by day 14 of the challenge (Fig. 4G). Significantly lower ZIKV RNA was seen in the brains and female reproductive organs (Fig. 4H & I) of the MV-E2 and MV-E-NS1(6) mice than in the MV-NS1(2) mice and the PBS mice that succumbed to the challenge. The combination group had significantly lower viral RNA in the brains, and complete viral clearance in the female reproductive organs than the MV-E2 and MV-E-NS1(6) vaccinated animals. The viral clearance achieved by the combination vaccine MV-E2 & MV-NS1 can be correlated with it, eliciting the highest ZIKV neutralizing antibodies of all groups (Fig. 4E & J). Lower ZIKV neutralizing antibodies were observed in the MV-E2 and the MV-E-NS1(6) group, and, as expected, no neutralizing activity was seen in the MV-NS1(2) group. One animal from the combination vaccine and the MV-E-NS1(6) group died due to unrelated reasons on day 30.

### The combination vaccine, MV-E2 & MV-NS1(2), induces durable plasma cell responses

Three groups of five female *hCD46 IFNαβR*^-/-^ mice (10-to 12-weeks old) were immunized i.p. with the 2 × 10^5^ TCID_50_ of the combination vaccine, MV-E2 & MV-NS1(2) (i.e., 10^5^ TCID_50_ of each virus), rMV, or PBS on day 0 and boosted on day 21. Their bone marrow and spleen were harvested on day 49 and assessed for the presence of ZIKV-E-, ZIKV-NS1-, and MV-H-specific long-lived and short-lived plasma cells (Fig. 5A). High ZIKV-E and MV-H IgG responses were seen in the combination vaccine group, while ZIKV-NS1 IgG responses were variable within the group (Fig 5B-D). Induction of high ZIKV neutralizing titers was observed in the combination vaccine group (Fig.5F). ZIKV-E specific long-lived plasma cells (LLPCs) were detected in the bone marrow, and short-lived plasma cell (SLPCs) responses were detected in the spleen of the combination vaccine group (Fig. 5G). Similar to the low-level ZIKV-NS1 antibody responses, the ZIKV-NS1 LLPCs and SLPCs were also low-level (Fig. 5H). Additionally, the MV-H specific LLPCs and the SPLCs in the combination group were similar to those of the empty vector rMV group, indicating that the addition of ZIKV-E and NS1 did not affect the vector response (Fig. 5I).

**Fig. 5.**
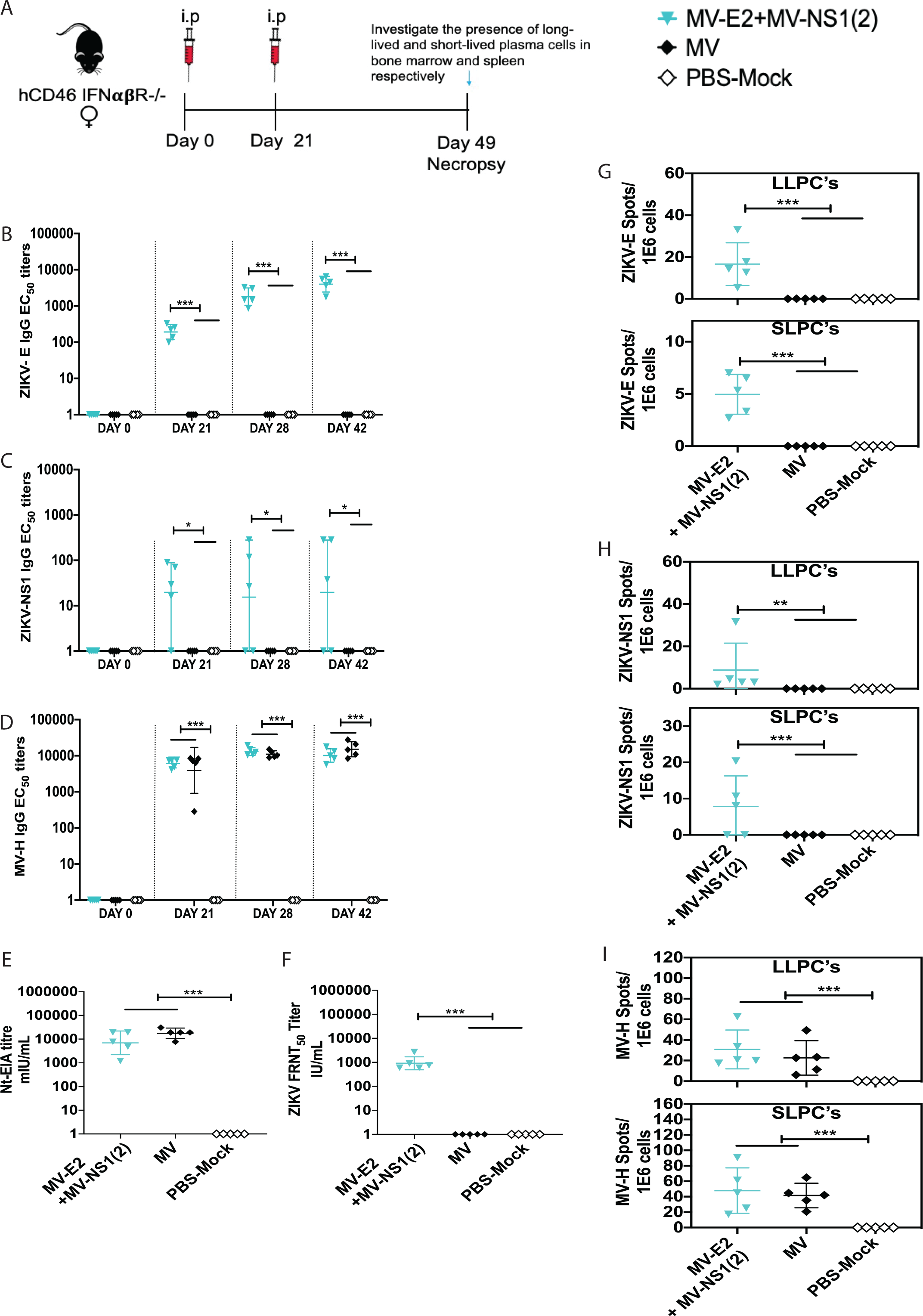
Long-lived and short-lived plasma cell response induced by the combination vaccine, MV-E2, and MV-NS1(2). **(A)** Timeline of vaccination, bone-marrow, and spleen harvesting. **(B-D)** anti-ZIKV-E (B), anti-ZIKV-NS1 (C), and anti-MV-H (D) ELISA EC_50_ titers of the vaccinated animals are plotted on a graph for all animals at different time points. The Mean ± SD is depicted per group. **(E)** MV Neutralization with low-passage MV Edmonston strain. Nt-EIA assay was performed on necropsy sera from vaccinated animals and controls. The 50% neutralizing titer (Nt-EIA titre) is plotted on the graph in terms of mIU/mL. The Mean ± SD is depicted per group. **(F)** ZIKV Neutralization with PRVABC59 Asian strain. FRNT was performed on necropsy sera from vaccinated animals and controls. The 50% neutralizing titer (FRNT_50_) is plotted on the graph in terms of IU/mL. The Mean ± SD is depicted per group. **(G-I)** ZIKV-E specific (G), ZIKV-NS1 specific (H), and MV-H specific (I) long-lived and short-lived plasma cell responses. The Mean ± SD is depicted per group. Statistics for Fig. 5B-I were done using the one-way ANOVA with posthoc Tukey HSD test and performed on log-transformed data for each time point. P-value of 0.12(ns), 0.033(*), 0.002(**), <0.001(***) are depicted accordingly.

### Measles virus neutralizing titers induced by the combination vaccine, MV-E2 & MV-NS1(2)

Sera from the previous experiment (Fig. 5) were assessed for the presence of MV-neutralizing titers. The combination vaccine group induced similar measles neutralizing titers to the empty vector rMV group (Fig. 5E), indicating that measles immunity was unaffected by the addition of ZIKV-E and NS1.

### Efficacy of the combination vaccine, MV-E2 & MV-NS1(2), in pregnant mice using a lethal mouse-adapted Zika African MR766 strain

Based on results indicating that the combination of MV-E2 and MV-NS1(2) can protect the female reproductive tract (Fig. 4), we conducted additional research using this combination vaccine in a pregnant mouse model. Three groups of ten female *hCD46 IFNαβR*^-/-^ mice (10-to 12-weeks old) were immunized i.p with the 2 × 10^5^ TCID_50_ of the combination vaccine, MV-E2 & MV-NS1(2) (i.e., 10^5^ TCID_50_ of each virus), and two PBS groups on day 0 and boosted on day 21 (Fig. 6A). The mice generated high ZIKV-E and MV-H antibody titers (Fig. 6B & D), but low-level to no ZIKV-NS1 antibodies (Fig. 6C) similar to the results of the previous experiments (Fig. 4C and 5C). Superovulation was induced in the mice by injecting 5 IU/mouse of pregnant mare serum gonadotropin (PMSG; i.p.) on day 36 (day-3 to pregnancy), and 5 IU/mouse of human chorionic gonadotropin (hCG; i.p.) on day 38 (day-1 to pregnancy). C57BL/6 males were placed in the cage, and the next day was considered day 0 of pregnancy. The mice were checked for plugs, weighed daily, and challenged with 10^2^ FFU of lethal ZIKV African MR766 strain on day 49-56, depending on pregnancy status being embryonic age 10.5-11.5 days, and humanely euthanized on day 17.5-18.5. The pregnant mice in the combination vaccine group had high ZIKV-E and moderate NS1 IgG responses (as shown by the blue filled black triangle). One PBS group was mock challenged with PBS. The combination vaccine group and the PBS group had three pregnant mice each, while the PBS-mock challenged group had four pregnant mice. The ZIKV-challenged PBS group showed signs of severe ZIKV disease at the necropsy timepoint, while the combination vaccinated animals showed no signs of disease. The pregnant females in the combination vaccine group had significantly lower or no viral RNA in the blood, brain, and placenta than the PBS group (Fig. 6F). Fetal heads from the combination vaccine group showed no presence of ZIKV RNA, with 88.5% intact fetuses and the remaining resorbed (Fig. 6G). The PBS challenged fetuses had significantly higher ZIKV RNA presence in the fetal heads, with 28% intact and 72% resorbed fetuses. Meanwhile, 94.7% intact and 5.3% resorbed fetuses were seen in the PBS unchallenged mice. The combination vaccine group had high ZIKV neutralizing titers on day 28 and at necropsy (Fig. 6E & H).

**Fig. 6.**
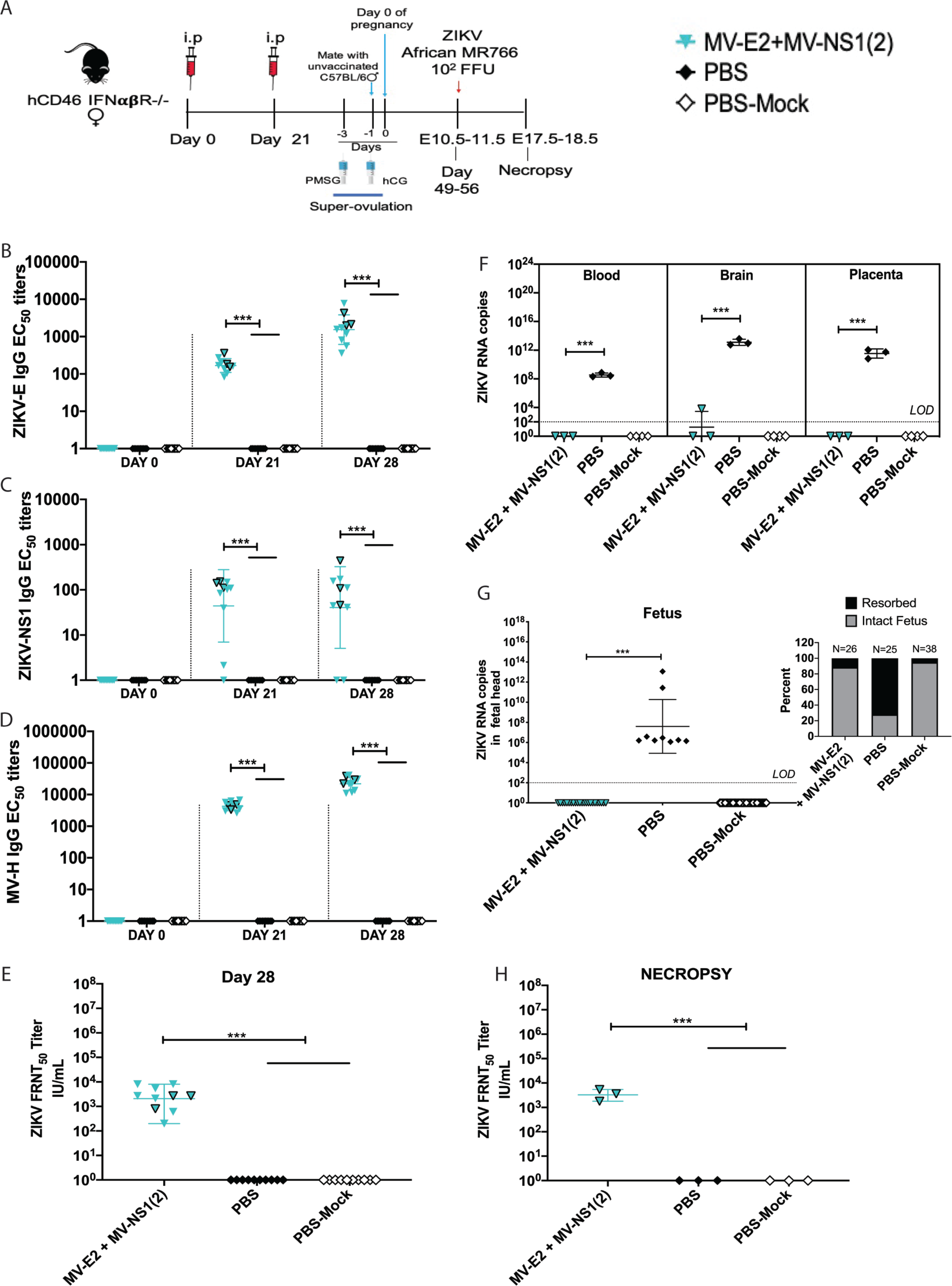
Immunogenicity and efficacy testing of the combination vaccine, MV-E2, and MV-NS1(2), using a lethal ZIKV African MR766, challenge strain in a pregnant mouse model. **(A)** Timeline of vaccination, super-ovulation, and challenge. The mice depicted as black-outlined blue inverted triangles(∇) are the pregnant mice in the combination group. **(B-D)** anti-ZIKV-E (B), anti-ZIKV-NS1 (C), and anti-MV-H (D) ELISA EC_50_ titers of the vaccinated animals are plotted on a graph for all animals at different time points. The Mean ± SD is depicted per group. **(E & H)** FRNT assay with PRVABC59 Asian strain was performed on day 28 (E) and Necropsy (H) sera. The 50% neutralizing titer (FRNT_50_) is plotted on the graph in terms of IU/mL. The Mean ± SD is depicted per group. **(F-G)** ZIKV RNA copies by qPCR in the blood, brain, and placenta of the pregnant mice (F) in the fetal head (G). Inset (G) percentage of resorbed and intact fetuses per group.The Mean ± SD is depicted per group. Statistics for Fig. 6E-H were done using the one-way ANOVA with posthoc Tukey HSD test and performed on log-transformed data for each time point. Only significant differences are depicted. P-value of 0.12(ns), 0.033(*), 0.002(**), <0.001(***) are depicted accordingly. LOD stands for limit of detection. A horizontal line () is used to include all groups below it.

## Discussion

Several candidate ZIKV vaccines are in Phase 1 clinical trials, with the most advanced being the DNA, mRNA, live-attenuated Zika virus (rZIKV/D4Δ30-713), and the alum adjuvanted Zika purified inactivated virus vaccines (*17*). Both the DNA vaccine and the inactivated virus vaccines are safe platforms that have been well-tolerated in humans (*18, 20*). Despite these results, recent studies with the same DNA vaccine found that it did not completely prevent adverse fetal outcomes in pregnant monkeys under prolonged ZIKV exposure (*25*). Another candidate vaccine—the Zika purified inactivated virus (ZPIV) vaccine failed to induce durable immune responses beyond eight weeks of vaccination in phase 1 clinical trials (*36*). This data suggests that further development in ZIKV vaccine strategies is paramount for the protection of susceptible pregnant women and their unborn fetuses. We tested the effectiveness of our combination MV-based ZIKV vaccine in the pregnant mouse model, and our findings indicated that a combination of MV-E2 and MV-NS1(2) vaccine could protect the female reproductive tract and their unborn fetuses.

There have been concerns that a ZIKV vaccine induces cross-reactive E protein-specific (mostly envelope domain I/II, fusion loop epitope-FLE) antibodies shared across flaviviruses and can potentiate antibody-dependent enhancement (ADE). ADE has been described among the dengue virus (DENV) strains and has had implications for the development of a DENV vaccine. Several *in-vitro* and *in-vivo* mouse studies have confirmed ADE of ZIKV with DENV antibodies and vice versa (*37*). ADE caused by the presence of DENV antibodies has been speculated to be one of the reasons for ZIKV-induced microcephaly (*38*). The hypothesis that transcytosis of IgG-virion complexes can occur across the placenta by utilizing the neonatal Fc receptor (FcRn) emerged from *in-vitro* studies in cytomegalovirus (CMV) and HIV (*39*). *In-vitro* studies of ZIKV enhancement in human placental tissue in the presence of DENV antibodies strengthened the hypothesis (*40*). However, human and monkey studies have indicated that prior DENV exposure provides some cross-protection to ZIKV infection, and vice versa (*41, 42*). Therefore, we do not expect our vaccine to cause ADE of DENV.

Different ZIKV strains have varied pathogenicity, with the Asian PRVABC59 strain being non-pathogenic and the African MR766 strain being extremely lethal in *IFNαβR*^-/-^ mice. The African MR766 strain is also the most potent at causing brain damage and postnatal lethality in mice (*43*). We investigated the immunogenicity and efficacy of MV vaccine vector-based ZIKV vaccines in *hCD46 IFNαβR*^-/-^ mice. The first-generation ZIKV vaccines, both expressing the full-length preME, MV-E0, and the MV-E2 vaccines, induced high ZIKV-E neutralizing antibodies and were protective when challenged with the non-lethal Asian PRVABC59 (10^6^ FFU) strain and the lethal African MR66 (10^4^ FFU) strain. Both vaccines achieved neutralizing titers above the correlate of protection determined by the adjuvanted purified inactivated Zika vaccine in rhesus macaques but did not achieve complete viral clearance in the lethal challenge model (*44*). The ZIKV-3′UTR-LAV-, GAd-Zvp-, and the E-dimer-vaccinated mice had similarly incomplete ZIKV clearance in organs when challenged with another ZIKV African strain (Dakar) (*24, 45-47*).

Recent studies have indicated that ZIKV NS1 antibodies and T cell responses play protective roles (*48*). But interestingly, we found that the NS1 antibodies themselves did not protect the mice from the lethal African MR766 challenge, as the MV-NS1(2) vaccinated animals succumbed to ZIKV disease. Similar results were observed in NS1-DNA vaccinated animals (*49*). Both the MV-E2 and MV-E-NS1(6) mice survived the challenge and showed significantly lower ZIKV RNA in the blood and brain of these mice compared to the PBS mice. We observed almost complete viral clearance in the female reproductive tract MV-E-NS1(6) vaccinated mice, while the combination group completely cleared ZIKV RNA from the reproductive tract, suggesting that NS1 antibodies played a role in this enhanced protection. The combination vaccine, MV-E2 & MV-NS1(2), also protected the fetuses in a lethal pregnant mouse model. In addition, the induction of ZIKV-E and ZIKV-NS1 long-lived plasma cell (LLPC) and short-lived plasma cell (SLPC) responses were seen in the combination vaccine group. Such durable responses have only been observed in the GAd-Zvp vaccinated animals that showed the presence of ZIKV-E LLPC’s and memory B cells (*45*). Taken together, the combination vaccine, MV-E2 & MV-NS1(2), is the first vaccine to provide complete fetal protection and viral clearance in the placenta when challenged with the African MR766 strain.

Past studies had shown that high E specific neutralizing titers were predictive of protection for other flaviviruses, however high ZIKV-E specific neutralizing were not sufficient to completely prevent adverse fetal outcomes in mice for several candidate vaccines (*24, 46*). Comparative studies of soluble preME (–Stem/TM) and full-length preME as the immunogen using either the DNA vaccine or gorilla adenovirus vector (GAd) or VSV vector indicated that the full-length prME might provide better protection in terms of controlling ZIKV replication (*45, 50-52*). The full-length preME assembles into a subviral particle (SVP) while the preME (–Stem/TM) is secreted as a soluble E protein alone, which may affect the epitopes exposed and thereby affect the quality of antibodies generated (*53*). The SVP characterization of the MV-ZIKV vaccines that express either prME or prME-NS1 suggests that it makes SVPs similar to the Zika virus, thereby exposing antigenically relevant epitopes to the immune system. While the NS1 expressed by MV-ZIKV vaccines are mostly a mixture of monomeric and dimeric forms, suggesting that the candidate vaccines make the membrane bound NS1. The secreted membrane bound NS1 may expose epitopes that allow for better targeting of infected cells.

Our findings add to the knowledge base on how a vaccine could be designed in order to provide complete fetal protection against ZIKV. Previously tested mRNA, DNA, and GAd-Zvp vaccines using the preME or preME-FLE (ZIKV-E fusion loop epitope mutant) as the antigen have been found to induce very high ZIKV neutralizing titers but failed to achieve complete placental and/or fetal protection in the vaccinated pregnant mice and primates (*24, 25, 45, 54*). An exception to this finding was demonstrated in the Jagger et al. study that observed complete placental and fetal protection in an mRNA-LNP vaccinated hSTA2-KI mouse model using the MA-ZIKV Dakar 41525 strain. This may be due to the reduced viremia induced by the MA-Dakar 41525 strain and the immunocompetent model allowing for adequate innate immune responses to control viral spread (*55*). ZIKV-E FLE antibodies have demonstrated ADE activity *in-vitro* and in mice and are suspected to be causing the adverse fetal outcomes. An alternative approach of using two DIII (domain III is the receptor-binding domain) monoclonals was investigated and found to reduce fetal pathology in primates but did not prevent maternal viral spread (*56*). Another group showed that a measles-based vaccine expressing preME (–Stem/TM) did not clear ZIKV from the placenta of the vaccinated animals in a low-dose Asian strain challenge, verifying the importance of full-length preME as the immunogen (*57*).

NS1 antibodies and T cell responses provided partial protection or protection in low-dose challenge models using the VSV and DNA vaccine platforms, but the MVA-NS1 vaccine was the only NS1 vaccine found to protect against a lethal African strain challenge in CD-1/ICR mice (*13, 23, 49, 51*). The two LAVs, 3□UTR-Δ10-LAV and the ZIKV-NS1-DKO (4 amino acid substitutions in the NS1 glycosylation sites), were tested in pregnant mouse models, and the 3□UTR-Δ10-LAV protected pregnant mice from the Asian PRVABC59 strain while the ZIKV-NS1-DKO failed to clear the ZIKV from fetal brains. Neither LAV study reported the NS1-specific antibodies or T cell responses induced by the vaccine (*24, 58*). In addition, the DNA vaccine that protected non-pregnant monkeys in an Asian PRVABC59 strain challenge failed to completely prevent maternal viremia and adverse fetal outcomes in pregnant monkeys (*15, 25*). Human studies on maternal antibodies of microcephalic newborns have observed an increase in ZIKV neutralizing capacity, with antibodies directed towards EDIII and lateral ridge EDIII antibodies in comparison to control newborns without microcephaly. This study also highlighted that mothers of microcephalic infants developed much lower NS1 antibodies than control newborns without microcephaly (*59*). Conversely, another study by the same group showed that antibody responses in individuals who developed high anti-ZIKV neutralizing antibodies had high EDIII and EDIII lateral ridge epitope antibodies (*60*). The two Robbiani et al. results congruently verify our hypothesis that both ZIKV-E and NS1 responses are needed for placental and fetal protection. To further corroborate this theory, high ZIKV NS1 antibody responses were observed in healthy Thai people who had high ZIKV neutralizing antibodies (*11*).

Collectively, our study and the published data suggest that ZIKV-E neutralizing antibodies protect non-pregnant mice and monkeys, while NS1 antibodies cannot provide such protection by themselves. An exception to this is the MVA-NS1 vaccine, which was tested in another mouse model. While ZIKV-E neutralizing antibodies are essential, the NS1 antibodies and T cell responses may aid its faster viral clearance (*49, 61*). More importantly, CD4^+^ T cells were recently identified as playing a vital role in the protection from ZIKV-induced neurologic disease and viral control (*62*). While the measles vaccine induces robust CD4^+^ and CD8^+^ T cell responses in infants (> 6 months) and adults, the CD4^+^ and CD8^+^ T cell responses are yet to be examined for the MV-ZIKA combination vaccine (*63*). Also, the low titer or incomplete seroconversion against NS1 induced by the combination vaccine may require further adjustment in the vaccine titers of the MV-E2 and MV-NS1(2) vaccines. Production and safety of the combination vaccine should follow the precedent of the measles vaccine, making it an ideal childhood vaccine platform. Hence, the combination vaccine, MV-E2 & MV-NS1(2), is a promising candidate ZIKV vaccine that warrants further preclinical development.

## Materials and Methods

### Experimental Design

Five *hCD46 IFNαβR*^-/-^ (IFNARCD46tg) breeding pairs were received from Dr. André Lieber (University of Washington, Seattle). Both male and female *hCD46 IFNαβR*^-/-^ mice were used in this study. For the pregnant mouse model, *hCD46 IFNαβR*^-/-^ mice were housed individually in microisolator cages. 8 to 10 weeks-old C57BL/6 male mice were purchased from Charles River for the pregnant mouse model.

Immunizations were conducted by inoculating mice with vaccines in 100 μl via intraperitoneal (i.p) or intramuscular route (i.m., 50 μl into each gastrocnemius muscle). ZIKV challenges were performed by subcutaneous inoculation in the hind limb with 10^4^ FFU of the mouse-adapted ZIKV African MR766 strain or 10^6^ FFU of the ZIKV Asian (PRVABC59 strain) in 100 μl PBS in non-pregnant mice. For the pregnancy experiments, super-ovulation was induced as the *hCD46 IFNαβR*^-/-^ female mice produce 5-6 pups under normal breeding conditions. *hCD46 IFNαβR*^-/-^ female mice were i.p. injected with 5IU/mouse of pregnant mare serum gonadotropin (PMSG) on day 36 and 5IU/mouse of human chorionic gonadotropin (hCG) on day 38 to induce super-ovulation. Super-ovulated *hCD46 IFNαβR*^-/-^ females were mated with naive wild-type C57BL/6 male mice on day 38. Females were checked for plugs the next day (day 39) and weighed daily until the end of the experiment. At E10.5-11.5, C57BL/6 male mice were removed from the cage and pregnant dams (*hCD46 IFNαβR*^-/-^) were inoculated with 10^2^ FFU of the mouse-adapted ZIKV African MR766 strain by subcutaneous injection in the hind limb. Animals were sacrificed at E17.5-18.5, and placentas, fetuses, and maternal tissues were harvested. All African challenged mice were euthanized when ethically defined clinical endpoints were reached (hind-limb paralysis). Mice were randomly allocated to groups. All experiments had five mice per group, except for the pregnant mouse study. For the pregnant mouse study-10 mice vaccinated, and 20 control animals were mated, wherein three vaccinated and three PBS mice and four PBS-mock challenged mice were impregnated.

### Animals and care

This study was carried out in strict adherence to recommendations described in the Guide for the Care and Use of Laboratory Animals (39), as well as guidelines of the National Institutes of Health, the Office of Animal Welfare, and the United States Department of Agriculture. All animal work was approved by the Institutional Animal Care and Use Committee (IACUC) at Thomas Jefferson University (animal protocol 01155 and 01873). All procedures were carried out under isoflurane anesthesia by trained personnel under the supervision of veterinary staff. Mice were housed in cages, in groups of five, under controlled conditions of humidity, temperature, and light (12-h light/12-h dark cycles). Food and water were available ad libitum.

### Cells

Vero-CCL81, Vero-E6, and 293T/T17 cells were purchased from ATCC and maintained in high glucose Dulbecco’s modified Eagle’s medium (DMEM, Corning, 10-017-CV) supplemented with 5% fetal bovine serum (FBS, R&D SYSTEMS, S11150) and 1% penicillin/streptomycin (P/S, Gibco, 15140122) and cultured at 37°C with 5% CO_2_.

### Viruses

The following ATCC viral stocks were purchased for this study. Zika virus African MR766 strain (ATCC, VR-1838), Zika virus Asian PRVABC59 strain (ATCC, VR-1843), and measles virus low-passage Edmonston strain (ATCC, VR-24).

### Antibodies

The following antibodies were used in this study: Anti-ZIKV-E mouse monoclonal antibody (Biofront Technologies, 1176-56), Pan-Flavivirus-E 4G2 mouse monoclonal antibody produced from hybridoma cell line D1-4G2-4-15 (ATCC, HB-112), Anti-Measles Nucleoprotein mouse monoclonal antibody produced from hybridoma NP.cl25 (Millipore Sigma, Cat # 95051114).

### Recombinant MV-ZIKV vaccines plasmid construction

To generate the first-generation MV-ZIKV vaccines the codon optimized Zika prME gene (signal peptide= MR766 strain, GenBank: MK105975.1 and prME sequence= PRVABC59 Asian 2015; GenBank: KU501215.1; Supplemental Fig. S1) was synthesized by GenScript. PCR amplification of the coding region of Zika prME from the gene synthesized plasmid was performed using primers ZMP Fwd1 (5’-GTGTCGACGCGTGGAATCCTCCCGTACGGCCACCATGGGGGCTGATA CAAGCATTGGCA - 3’) and ZMP Rev1 (5’-GTGTCGGACGTCATTTATGCGGACACTGCG GTGGACAGAAAA-3’). PCR fragments were digested with the respective enzymes and ligated into the ATS-0 or ATS-2 MV vector to generate MV-E0 and MV-E2, respectively (JM109 *E*.*coli*). The MV virus (Edmonston B strain) vaccine vector expressing GFP at ATS-0 flanked by BsiWI, and AatII restriction sites were received from Dr. R. Cattaneo. This vector was modified to include a hammerhead ribozyme at the 5’ end (ATS-0 MV vector). A MV (Edmonston B strain) vaccine vector (ATS-2 MV vector) was generated to create an additional transcriptional site (ATS) by inserting MluI and AatII sites at ATS-2.

To generate the 2^nd^ generation MV-ZIKV vaccines, the codon optimized Zika prME-NS1 gene (strain PRVABC59 Asian 2015; GenBank: KU501215.1; Supplemental Fig. S5) was synthesized by GenScript. PCR amplification of the coding region of Zika prME-NS1 from the gene synthesized plasmid was performed using primers MV-ZprME-NS1(2) Fwd (5’-ACAGAGTGATACGCGTACGGGCCACCATGGGGG-3’) and MV-ZprME-NS1(2) Rev (5’-GCACGCGATCGCAAGACGTCGGCTATGCTGTCACC-3’). PCR fragment was inserted into the ATS-2 MV vector to generate MV-E-NS1(2) by In-Fusion cloning (Stellar cells). An additional MV (Edmonston B strain) vaccine vector (ATS-6 MV vector) was generated to create an additional transcriptional site (ATS) by inserting MluI and SpeI sites at ATS-6. PCR amplification of the coding region of Zika prME-NS1 from the gene synthesized plasmid was performed using primers MV-coZprME-NS1(6) Fwd (5’-ACAGAGTGATACGCGTACGGGCCACCATGGGGGCTGA TAC-3’) and MV-coZprME-NS1(6) Rev (5’-TCTATTTCACACTAGTGCGATCGCGACGTCG GCTATGCTGTCACC-3’). The PCR fragment was digested and inserted into ATS-6 MV vector to generate MV-E-NS1(6). PCR amplification of the coding region of Zika prME-NS1 from the gene synthesized plasmid (Supplemental Fig. S6) was performed using primers MV-coNS1 Fwd 1 (5’-ACAGAGTGATACGCGTACGGGCCACCATGGAGACAGACACACTCCT-3’) and MV-coNS1 Rev 1 (5’-GCACGCGATCGCAAGACGTCGGCTATGCTGTCACCATAGAGCG GACCAGGTTG-3’) was used to generate fragment 2. Fragment 2 was digested by restriction enzymes MluI, SgrAI. In order to insert the human IgKappa signal peptide at ATS-2 to allow for better secretion of ZIKV-NS1, fragment 1 was generated by PCR amplification using primers IgKappa Fwd 2 (5’-ACAGAGTGATACGCGTGGCCACCATGGAGACAGACACACTCCTGC TATGGGTACTGCTGCTCTGGGTTCCAGGTTCCACG-3’) and IgKappa Rev 2 (5’-ACACGAACACGCCGGTGCCACACCGTGTCTCCTTCTTAGAAAAATCCACGGAGCATCCCACGTCACCAGTGAACCTGGAACCCAGA-3’). The PCR fragments 1, and 2 were inserted into the ATS-2 MV vector to generate MV-NS1(2) by In-Fusion cloning (Stellar cells).

### Virus recovery

293T/T17 cells (0.8 ×10^6^ cells per well) were pre-seeded in 6 well plates. The MV-ZIKV viruses and control MV’s were rescued from their full-length cDNA with the helper-plasmid rescue system. 293T/T17 cells were transfected with pCAGGS-T7, pTIT-MVN (MV-Nucleoprotein), pTIT-MVP (MV-Phosphoprotein), and pEMC-MVLa (MV-Large protein), using the XtremeGene9 reagent (Millipore Sigma). After overnight incubation at 37°C, the cells were heat shocked at 42°C for 3 hours and then returned to 37°C. After 3 days of incubation at 37°C, transfected cells were transferred onto a monolayer of Vero cells and incubated at 37°C. Virus was harvested from Vero cells when syncytia involved 80% to 90% of the culture by scraping infected cells, freeze-thawing cells, and medium, and centrifuging them to remove cellular debris.

### Virus production

The Measles viruses were grown in T175 flask with sub-confluent Vero cells in Optipro SFM. The medium was changed every 3-4 days, and supernatant was collected until the cell monolayer came-off. The low passage MV Edmonston strain (ATCC-VR 24) was propagated by inoculating 100 μL of the original ATCC vial into a T175 flask pre-seeded with Vero cells (80% confluent) in 32 mL of 1X DMEM (2% FBS) and placed 37°C in a humidified 5% CO_2_ atmosphere. The virus supernatant was harvested after 12 days. The virus was harvested by freeze-thawing cells and medium, centrifuging them at 3000 rpm for 10 minutes to remove cellular debris. Viral supernatants were tittered, aliquoted, and frozen at –80°C.

Zika viruses were grown in a T175 flask with sub-confluent Vero cells in 1X DMEM (2% FBS, 1% PS). The medium was changed every 3-4 days, and the supernatant was collected until the cell monolayer came-off. The harvested virus was tittered on Vero cells, and high titer stocks were aliquoted and frozen at –80°C

### Virus titration

#### Measles virus titration

Measles virus titers were analyzed as 50% tissue culture infectious dose (TCID50) by the Reed–Muench method. 10^4^ Vero cells in 100 μL per well were pre-seeded in a 96 well flat-bottom plate 2 hours before the addition of the virus to the well. 180μL of 1X DMEM (Corning, Cat# 10-017-CV) was added to every well of the 96 well round bottom plate (dilution plate). 20 μL of one virus was added to Column 1 of the dilution plate. Twelve 10-fold serial dilutions of the virus were performed in the dilution plate in a total volume of 200 μL per well. 30μL of the diluted virus was transferred from one row to each row of the 96 well plates pre-seeded with cells, changing tips between each row. Two plates were prepared per inoculum. Plates were incubated at 37°C for 4 days. On day 4, plates were fixed with 80% Acetone for 10 mins at 4°C. The fixation solution was aspirated, and plates were allowed to air dry. Cells were blocked for 1hour in FACS buffer and stained with 100μL of Anti MV-N cl25 mouse monoclonal antibody (2 μg/mL in FACS buffer-1X PBS, 10%FBS, 0.05% Sodium azide) for 2 hours. Plates were washed 3X with FACS buffer and stained with secondary antibody, Cy3-conjugated Goat Anti-mouse IgG (2 μg/mL in FACS buffer) for 2 hours. After 3X washes with FACS buffer and plates were read using a fluorescence microscope. The TCID50 titer was calculated with the following formula:

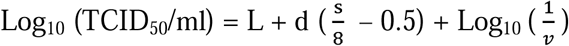

Where L is the reciprocal of the last dilution in which all well is positive, d is the log_10_ of dilution factor, v is the volume of inoculum (ml/well).

#### Zika virus titration

ZIKV stocks were propagated in Vero cells or C6/36 cells and titrated by focus-forming assay (FFA) as described previously (*64*). Briefly, ten-fold serial dilutions of ZIKV in 1X DMEM (Corning, Cat# 10-017-CV) supplemented with 2% fetal bovine serum (R&D systems, Cat# S11150) and 20U/mL Penicillin-Streptomycin (Gibco, Cat# 1540122) were performed in 96-well Costar (Corning, NY) plates. 2.5× 10^4^ Vero or C6/36 cells per well were added to the 96 well plate incubated undisturbed for 3 days at 37°C. Media overlay was aspirated, and the cell monolayer was fixed with 80% Acetone for 10 minutes at 4°C. The fixation solution was aspirated, and plates were allowed to air dry. Cells were blocked for 1hour in FACS buffer and stained with Pan-Flavivirus E mouse monoclonal (1 μg/mL in FACS buffer) for 2 hours. Cells were then washed 3X with FACS buffer and stained with secondary antibody, Cy3-conjugated Goat Anti-mouse IgG (Jackson ImmunoResearch, 2 μg/mL in FACS buffer) for 2 hours. After 3X washes with FACS buffer and plates were read using a fluorescence microscope. For each sample, a dilution with easily distinguished foci is selected, and titer is calculated in focus-forming units per ml (FFU/ml), using the average of triplicate wells:

### Multi-step growth curve

We infected Vero-CCL81 cells with wt or recombinant viruses at a multiplicity of infection (MOI) of 0.001, in triplicate. The infected cells and supernatant were harvested at 12, 24, 48, 72, 96, 120, and 144 hpi, respectively. After three cycles of freeze-thawing and sonication of infected cells, lysates were centrifuged to remove cellular debris, and the supernatant was collected. The titer (TCID_50_/ml) of each sample was measured using Vero-CCL81 cells.

### Immunofluorescence assay (IFA)

8.5 × 10^4^ Vero-E6 cells were seeded in 24 well plates with coverslips. After 18 hours, the cells were infected with the recovered MV-ZIKV viruses and controls at a MOI of 0.1 for 72 hours. The cells were permeabilized for 20 minutes at room temperature with BD Cytofix/Cytoperm (BD, 554714) and blocked with FACS buffer for 30 minutes. For Fig. 1B, Cells were stained with Biofront ZIKV-E mouse monoclonal antibody (1μg/mL) for 1 hour at room temperature (RT) on a rocker platform. Cells were washed 3X with FACS buffer and stained with secondary antibody Cy3-conjugated Goat Anti-mouse IgG (Jackson ImmunoResearch, 2 μg/mL in FACS buffer) for 1 hour at RT. For Fig. 3B, cells were stained with Biofront ZIKV-E mouse monoclonal antibody (1 μg/mL) and Anti-ZIKV NS1 human monoclonal antibody EB9 (2μg/mL) for 1 hour at room temperature (RT) on a rocker platform. Cells were washed 3X with FACS buffer and stained with secondary antibodies Alexa Fluor 568 -conjugated goat anti-mouse IgG (ThermoFisher Scientific, 2.5 μg/mL), and Alexa Fluor 647 conjugated goat α-human IgG (ThermoFisher Scientific, 2.5 μg/mL), for 1 hour at RT. For both Fig. 1B & 4B, cells were washed 3X with FACS buffer and stained with Anti MV-N cl25 mouse monoclonal antibody conjugated with Dylight 488 (5 μg /mL) at RT for 1 hour. Cells were washed 3X with FACS buffer and mounted with VECTASHIELD® Hardset™ Antifade Mounting Medium with DAPI (H-1500). Images were taken using Nikon A1R+ confocal microscope.

### Viral sucrose purification and Cell lysates

Larger amounts of MV-ZIKV and control MV supernatants were spun through a 20% sucrose cushion in an SW32 Ti rotor (Beckman, Inc.) at 25,000 rpm for 2 hours. ZIKV was spun through a 20% sucrose cushion at 30,000 rpm for 3.5 hours. Virion pellets were resuspended in phosphate-buffered saline (PBS), and protein concentrations were determined using a bicinchoninic acid (BCA) assay kit (Pierce). 6 well plates seeded with 0.7 × 10^6^ Vero cells, 16 hours before they were infected with MV-ZIKV viruses and control viruses at a MOI of 5 for 60 hours and harvested using Sabatini Buffer (40mM Tris,ph7.6; 120 mM NaCl; 1mM TRITON-X100; 0.4mM Sodium Deoxycholate; 1mM EDTA), and protein concentrations were determined using a bicinchoninic acid (BCA) assay kit (Pierce).

### Purification of Subviral particles (SVPs)

T175 flasks were infected with MV-ZIKV vaccines or controls at a MOI of 0.1. Larger amounts of MV-ZIKV and control MV and Zika virus supernatants were filtered through a 0.2 μm filter (Rapid-Flow™ Sterile Disposable Bottle Top Filters with PES Membrane). The 0.2 μm filtration step was aimed to filter most of the MV particles (pleomorphic, 100-300 nm) out of the supernatant. The filtered supernatant was then spun through a 20% sucrose cushion at 48,000 rpm for 3 hours in a SW55Ti rotor (Beckman Coulter). The SVP pellets were resuspended in a 4X non-reducing buffer (Alfa Aesar, ThermoFisher, Cat# J63615-AD) in a total volume of 100μL. SVPs for MV-E0 virus could not be purified and were therefore not included in this blot.

### SDS-PAGE and Western Blot

The sucrose purified virus particles or cell lysates were denatured with 4X Laemmli Sample Buffer supplemented with 2-mercaptoethanol (10%) at 95°C for 10 minutes. 2.5 μg of sucrose purified virus or 5 μg of cell lysates or 10μL of the SVPs (in non-reducing buffer) was resolved on a 10% SDS–polyacrylamide gel and thereafter stained overnight with SYPRO Ruby for total protein analysis or transferred onto a nitrocellulose membrane in Towbin buffer (192 mM glycine, 25 mm Tris, 20% methanol) for Western blot analysis. The nitrocellulose membrane was then blocked in PBST (1X PBS, 0.05% Tween-20) containing 5% dried milk at room temperature for 1 hour. After blocking, the membrane was washed 3X with PBST and incubated overnight with Biofront ZIKV-E mouse monoclonal antibody (1μg/mL) or Anti MV-H Rabbit polyclonal sera (diluted 1:5000) or Anti MV-N cl25 mouse monoclonal antibody (1μg/mL) or Anti ZIKV-NS1 B4 mouse monoclonal antibody (1μg/mL) in 10% bovine serum albumin (BSA). After washing, the blot was incubated for 1 h with horseradish peroxidase (HRP)-conjugated anti-mouse/human/rabbit IgG diluted 1:20,000 in blocking buffer depending on the primary antibody used. Bands were developed with SuperSignal West Dura Extended duration substrate (Pierce).

### ZIKV Envelope Antigen

Recombinant Zika Envelope protein antigen: The antigen used for ELISA and ELISPOT assay was purchased from Aalto BioRegents (AZ 6312).

### Recombinant Measles virus H and ZIKV-NS1 antigen

A codon optimized MV-H gene (Edmonston B strain) was gene synthesized by GenScript. PCR amplification of the coding region of MV-H from the plasmid was performed using primers coMV-H61 N HA FWD (5’-TCGTGGTGCCAGATCTCACAGAGCCGCCATCTAT - 3’) and coMV-H61 N-HA REV (5’-TCTCGAGCGGCGGCCGCCTACCTTCTATTTGTGCCG -3’) to generate a fragment from amino acid 61 to 617 of MV-H protein (Supplemental Fig S9). The PCR amplified fragment was inserted by In-Fusion cloning (Stellar cells) into the pDisplay vector containing N terminal hemagglutinin (HA) tag that was cut with restriction enzymes BglII and NotI.

A codon optimized Zika prME-NS1 gene (strain PRVABC59 Asian 2015; GenBank: ANW07476.1) was gene synthesized by GenScript. The recombinant ZIKV-NS1 was constructed as published previously (*13*). Briefly, PCR amplification of the coding region of ZIKV-NS1 from the plasmid was performed using primers Sol coNS1 Fwd (5’-TGACGCACCTAGATCTAATGG CTCCATCTCTCTGATGTGC -3’) and Sol coNS1 Rev1 (5’-CGTATGGATAGTCGACAGCACGTCCTGCTGTCACCATAGAGCGGACC-3’) to generate a fragment that incorporated the last 24 amino acids of ZIKV envelope (NGSISLMCLALGGVLIFLSTAVSA) to the amino terminus of the NS1 coding region (Amino acid 1-352). The PCR amplified fragment was inserted by In-Fusion cloning (Stellar cells) into the pDisplay vector containing C terminal HA tag that was cut with restriction enzymes BglII and SalI.

Sub-confluent T175 flasks of 293T cells (human kidney cell line) were transfected with XtremeGene9 (150 μL/flask) and a pDisplay vector encoding either the codon optimized MV-H fused to an N-terminal HA tag (50μg/flask) or codon optimized ZIKV-NS1 fused to an C-terminal HA tag (50μg/flask). Supernatant was collected between days 7 post transfection and loaded onto two different equilibrated anti-HA agarose (Pierce) columns containing a 2.5-ml agarose bed volume. The supernatant is recirculated overnight at 4°C using a peristaltic pump at 1ml/minute. The column was washed with 10 bed volumes of PBS (0.05% Sodium Azide). After washing, antibody-bound MV-H was eluted with 5 ml of 250 μg/ml HA peptide in PBS. Fractions were collected and analyzed for MV-H by Western blotting with monoclonal anti-HA-7 antibody (Sigma) prepared in 10% BSA. Peak fractions were then pooled and dialyzed against PBS in 10,000 molecular weight cutoff (MWCO) dialysis cassettes (Thermo Scientific) to remove excess HA peptide. After dialysis, the protein was quantitated by UV spectrophotometry and frozen in small aliquots at −80°C.

### Enzyme-linked immunosorbent assay

We tested individual mouse sera by enzyme-linked immunosorbent assay (ELISA) for the presence of IgG specific to ZIKV-E, ZIKV-NS1, and MV-H. To test for anti-ZIKV-E humoral responses, recombinant ZIKV-E (Alto BioRegents) was resuspended in a coating buffer (50 mM Na2CO3 [pH 9.6]) at a concentration of 1 μg /ml and then plated in 96-well ELISA MaxiSorp plates (Nunc) at 100 μl in each well. ZIKV-NS1 and MV-H were similarly resuspended in coating buffer (50 mM Na2CO3 [pH 9.6]) at a concentration of 1 μg/ml and then plated in 96-well ELISA MaxiSorp plates (Nunc) at 100 μl per well. After overnight incubation at 4°C, plates were washed three times with 1X PBST (0.05% Tween 20 in 1× PBS), which was followed by the addition of 250 μl blocking buffer (5% dry milk powder in 1× PBST) and incubation at room temperature for 1 hour. The plates were then washed three times with PBST and incubated overnight at 4°C with serial dilutions of sera in 1X PBST containing 0.5% BSA, 0.05% Sodium azide. Plates were washed three times the next day, followed by the addition of horseradish peroxidase-conjugated goat anti-mouse-IgG Fc secondary antibody (1:2000) (Southern Biotech, 1033-05). After incubation for 2 hours at room temperature, plates were washed three times with PBST, and 200 μl of o-phenylenediamine dihydrochloride (OPD) substrate (Sigma) was added to each well. The reaction was stopped by the addition of 50 μl of 3M H_2_SO_4_ per well. Optical density was measured at 490 nm (OD490) using BioTek Spectrophotometer. ELISA data were analyzed with GraphPad Prism 8. using a sigmoidal nonlinear fit model to determine the 50% effective concentration [EC50] titer. The EC50 titer is the concentration (dilution) at which the antibody/serum at which you get 50% of your maximal effect (Optical Density).

### Zika neutralization assay

A FRNT measured ZIKV neutralizing antibody was performed as previously described (*65*). Briefly, heat-inactivated (56°C, 30minutes) sera were serially diluted (three-fold) starting at a 1/30 dilution and incubated with 10^2^ FFU of ZIKV (strain /PRVABC59/2015/P1 Vero) for 1 hour at 37°C. The ZIKV-serum mixtures were added to Vero cell monolayers in 96-well plates (1.2 × 10^4^ Vero cells per well were seeded 16 hours prior to virus addition) and incubated for 1.5 h at 37°C, followed by overlaying the cells with 1% (w/v) methylcellulose in 1X DMEM (5%FBS). Cells were incubated for 40 hours at 37°C and subsequently fixed using 2% PFA in PBS for 1 hour at room temperature. Cells were permeabilized with Perm buffer (1X PBS, 5% FBS, 0.2% Triton X-100) for 20 minutes at 4°C and washed 3X with FACS buffer (1XPBS, 5% FBS, 0.05% Sodium azide). ZIKV-infected cell foci were detected using anti-Flavivirus E 4G2 mouse monoclonal

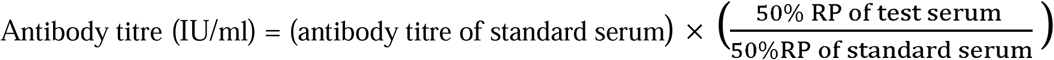

antibody (1ug/mL), washed 3X with FACS buffer, followed by Cy3-conjugated Goat Anti-mouse IgG (Jackson ImmunoResearch, 2 μg/mL). After 3X washes with FACS buffer and plates were read using a fluorescence microscope. For Fig.s 5, 6, and 7, the 1st International Standard for anti-Asian lineage Zika virus antibody (NIBSC: 16/352) and Working reagent for anti-Zika virus antibody (NIBSC: 16/320) were used at a starting dilution of 1:100. The 50% reduction point (RP -Reciprocal of the dilution where 50% neutralization is observed) for the serum and standards were noted, and the FRNT50 titer was calculated as follows:

### Measles neutralization assay

Measles neutralization assay was performed as described previously (*66*). Sera and the 3rd International standard for Anti-Measles serum were heat-inactivated at 56°C for 30 minutes. In a 96 well plate, serum samples were diluted serially 4-fold from 1/10, and the 3rd International standard was diluted 1/100 in 1X DMEM (2% FBS, 1%PS) 70 μl and mixed with 30 μl volume of diluted virus solution (150 PFU/well) of low-passage MV Edmonston strain (P1, Vero) in 1X DMEM (2% FBS, 1%PS) on a plate shaker at 37°C for 1 h. Then, 1.2×10^4^ Vero cell suspension was added (100 μl) and incubated for 68 hours at 37°C. Cells were fixed with 80% acetone for

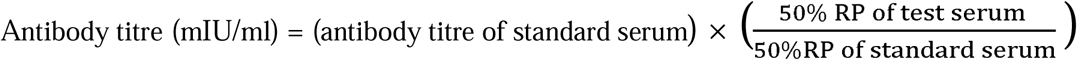

10 mins at 4°C. The fixation solution was aspirated, and plates were allowed to air dry. Cells were blocked for 1 hour in FACS buffer and stained with 100μL of Anti MV-N cl25 mouse monoclonal antibody (2 μg/mL in FACS buffer-1X PBS, 10%FBS, 0.05% sodium azide) for 2 hours. Plates were washed 3X with FACS buffer and stained with secondary antibody, Cy3-conjugated Goat Anti-mouse IgG (2 μg/mL in FACS buffer) for 2 hours. After 3X washes with FACS buffer and plates were read using a fluorescence microscope, the presence of the measles virus was detected by direct EIA as described above. All serum dilutions were tested in triplicate. The 50% reduction point (50%RP) of each serum was calculated using the Reed–Muench formula. The neutralizing antibody titer of test sera was converted into mIU/ml by comparing their 50% RP with that of the international standard serum using the following formula:

### RNA extraction

Whole blood (50 µL) was resuspended in 150 µL of TRIzol LS Reagent (Life Technologies). All Organs were added to Omni pre-filled bead tubes containing 1mL of TRIzol and homogenized using the OMNI bead ruptor 12. The RNA extraction protocol for biological fluids using TRIzol LS Reagent was followed until the phase separation step. The remaining RNA extraction was done using the PureLink RNA Mini Kit (Ambion). The quantity and quality (260/280 ratios) of RNA extracted was measured using NanoDrop (Fisher).

### Quantification of Zika virus RNA by quantitative Real-Time polymerase chain reaction

ZIKV cDNA was generated from RNA isolated from the ZIKV African MR766 strain and Asian PRVABC59 strain by One-Step RT PCR (SuperScript III, Thermo Fisher Scientific) with primers ZKV NS4B IVT F1 (5’-GAATTCTAATACGACTCACTATAGGGGCATCTAATGGGAAGG AGA-3’) and ZKV NS4B IVT R1 (5’-GCTAGCGGCTGTAGAGGAGTTCCAGTA-3’). The African and Asian standards were generated by *in-vitro* transcription of the generated ZIKV cDNA, followed by using the MEGAclear Transcription Clean-Up Kit. Aliquots of 2 ×10^10^ copies/µL were frozen at –80°C. Five microliters of RNA per sample were run in triplicate, using the ZIKV-F2 (5’-CAGCTGGCATCATGAAGAATC-3’) and ZIKV-R1 (5’-CACTTGTCCCATC TTCTTCTCC-3’) primers for African strain detection (ThermoFisher SCIENTIFIC) or the ZIKV-F1 (5’-CAGCTGGCATCATGAAGAACC-3’) and ZIKV-R2 (5’-CACCTGTCCCATCTTTTTC TCC-3’) primers for Asian strain detection. The panZika-Probe (6FAMGTTGTGGATGGAATA GTGGMGBFNQ) detects both the Asian and the African strain.

### ELISPOT

An ELISPOT assay quantitated the number of ZIKV E, ZIKV-NS1, and MV-H specific LLPCs and SLPCs in the bone marrow and spleen, respectively, as previously described (*67*). ELISPOT plates (Millipore) were coated with ZIKV-E antigen (50 μg/mL), ZIKV-NS1 antigen (50 μg/mL) and MV-H antigen (10 μg/mL), overnight at 4°C. Subsequently, plates were washed six times with PBS (200 μl) and then blocked for 1-2 hours with Goat Serum (ThermoFisher Scientific, 16210072) at 37°C. Bone marrow cells from the femurs and splenocytes were harvested from the immunized mice and controls, and erythrocytes were lysed by ACK lysis buffer. Subsequently, 3 ×10^6^ cells/well were added to the ZIKV-E, and ZIKV-NS1 coated plates, and 1 ×10^6^ cells/well were added to the MV-H coated plates. Cells were serially diluted in a 96 well round bottom plate, transferred to the coated ELISPOT plate, and incubated overnight at 37°C in a CO_2_ incubator for 16 hours. Plates were washed four times with 1X PBST, incubated with HRP conjugated goat anti-mouse IgG-Fc (Sothern Biotech, 1μg/mL) in PBS-T for 2 hours at 37°C. Following 3X washing with 1X PBST, plates were washed 3X with PBS. Spots were developed with TrueBlue peroxidase substrate (KPL) before the reaction was quenched with water and counted with an AID EliSpot Reader (Autoimmun Diagnostika GmbH).

### Statistical analysis

Specific statistical tests used to analyze experimental datasets are described in the respective Fig. Legends. For experiments with only female mice, antibody responses, and immune cell analyses, One-way ANOVA with posthoc Tukey HSD test was performed on log-transformed data for each time point. For data analysis where only two groups were compared, a Mann-Whitney U test was performed on log-transformed data for each time point. For experiments with female and male mice, antibody responses, and immune cell analyses, two-way ANOVA with posthoc Tukey HSD test was performed on log-transformed data for each time point. Survival curves were analyzed using the log-rank test with a Bonferroni correction. A P value of < 0.05 was assigned to establish statistical significance using GraphPad Prism version 8.0.

## General

We thank Jennifer Wilson (Thomas Jefferson University, Philadelphia, PA) for critical reading and editing the manuscript; Dr. Christian Pfaller and Dr. Robberto Cattaneo for providing the MV-GFP0 plasmid, measles H antibody, and measles titration protocols; Dr. Glenn Rall for providing the measles helper plasmids; Dr. André Lieber for transferring the *hCD46 IFNαβR*^-/-^ mice; Dr. Gene Tan for providing the human NS1 antibody; and Annamarie Testa for assisting in some experimental work.

## Author Contributions

D.K and M.J.S. designed the studies and wrote the paper. D.K performed and analyzed all experiments. C.W provided the modified measles vaccine vector, the measles GFP-Nluciferase2 plasmid, and the MV-E-NS1(6) plasmid.

## Data and materials availability

All data generated from this study are present in the paper or Supplementary Materials. Vaccines and Reagents can be shared with a standard material transfer agreement upon request to the corresponding author, Matthias J. Schnell.

## Supplementary Materials

**Fig. S1. Complete nucleotide and translation sequence of the ZIKV prME antigen**.

**Fig. S2. Immunogenicity and efficacy testing of first-generation candidate MV-ZIKV vaccines using a non-lethal ZIKV Asian PRVABC59 challenge strain**.

**(A)** Timeline of vaccination, challenge, and viral load determinations.

**(B-C)** Anti-ZIKV-E (B) and anti-MV-H (C) specific ELISA EC_50_ titers of the vaccinated animals are plotted on a graph for all animals at different time points. The Mean ± SD is depicted per group.

**(D & H)** ZIKV neutralization with PRVABC59 Asian strain. FRNT assay was performed on day 63 (D) and necropsy (H) sera from vaccinated animals and controls. The 50% neutralizing titer (FRNT_50_) is plotted on the graph. The Mean ± SD is depicted per group.

**(E-G)** ZIKV RNA copies by qPCR in the blood (E), brain (F), and reproductive tract (G). The Mean ± SD is depicted per group.

Statistics for Fig. S1B-C was done using the Mann-Whitney U test and performed on log-transformed data for each time point. Statistics for Fig. S1D-H was done using the one-way ANOVA with a posthoc Tukey HSD test an performed on log-transformed data for each time point. Only significant differences are depicted. P-value of 0.12(ns), 0.033(*), 0.002(**), <0.001(***) are depicted accordingly. LOD stands for limit of detection. A horizontal line () is used to include all groups below it.

**Fig. S3. Long-term immunogenicity and efficacy testing of first-generation candidate MV-ZIKV vaccines using a lethal ZIKV African MR766 challenge strain**.

**(A)** Timeline of vaccination, challenge, and viral load determinations.

**(B-C)** Anti-ZIKV-E (B) and anti-MV-H (C) specific ELISA EC_50_ titers of the vaccinated animals are plotted on a graph for all animals at different time points. The Mean ± SD is depicted per group.

**(D & I)** FRNT assay was performed on day 144 (D) and necropsy (I) sera from vaccinated animals and controls. The 50% neutralizing titer (FRNT_50_) is plotted on the graph. The Mean ± SD is depicted per group.

**(E)** Kaplan-Meier survival curve analysis of vaccinated and control animals post challenge.

**(F-H)** ZIKV RNA copies by quantitative polymerase chain reaction (qPCR) in the blood (F), brain (G), and reproductive tract (H). The Mean± SD is depicted per group.

Statistics for Fig. S2: B-D & F-I were done using the one-way ANOVA with posthoc Tukey HSD test and performed on log-transformed data for each time point. Fig. S2E survival curves were analyzed using the log-rank test with a Bonferroni correction. Only significant differences are depicted. P-value of 0.12(ns), 0.033(*), 0.002(**), <0.001(***) are depicted accordingly. A horizontal line () is used to include all groups below it.

**Fig. S4. Effect of intramuscular route of vaccination and prior MV immunity to the immunogenicity and efficacy of first-generation candidate MV-ZIKV vaccines using a lethal ZIKV African MR766 challenge strain**.

**(A)** Timeline of vaccination, challenge, and viral load determinations. The black boxed open pink triangle Δ is the MV-E0(M) animal that succumbed to ZIKV disease.

**(B-C)** Anti-ZIKV-E (B) and anti-MV-H (C) specific ELISA EC_50_ titers of the vaccinated animals are plotted on a graph for all animals at different time points. The Mean ± SD is depicted per group.

**(D & I)** FRNT assay was performed on day 63 (D) and necropsy (I) sera. The 50% neutralizing titer (FRNT_50_) is plotted on the graph. The Mean ± SD is depicted per group.

**(E)** Kaplan-Meier survival curve analysis of vaccinated and control animals post challenge.

**(F-H)** ZIKV RNA copies by qPCR in the blood (F), brain (G), and reproductive tract (H) at necropsy. The Mean± SD is depicted per group.

Statistics for Fig. S3: B-D & F-I were done using the two-way ANOVA with posthoc Tukey HSD test and performed on log-transformed data for each time point. Fig. S3E survival curves were analyzed using the log-rank test with a Bonferroni correction. Only significant differences are depicted. P-value of 0.12(ns), 0.033(*), 0.002(**), <0.001(***) are depicted accordingly. A horizontal line () is used to include all groups below it.

**Fig. S5. Complete nucleotide and translation sequence of ZIKV prME-NS1 antigen**.

**Fig. S6. Complete nucleotide and translation sequence of ZIKV NS1 antigen**.

**Fig. S7. Multi-step growth curves**.

**Fig S8: Subviral particle characterization**

**(A)** Western blot of sucrose purified SVPs probed for ZIKV-E

**(B)** Western blot of sucrose purified SVPs probed for ZIKV-NS1

**Fig. S9. Complete nucleotide and translation sequence of codon optimized MV-H (Edmonston B strain) antigen**.

## References

1. G. Calvet et al., Detection and sequencing of Zika virus from amniotic fluid of fetuses with microcephaly in Brazil: a case study. Lancet Infect Dis 16, 653–660 (2016).

2. B. Valle Borrego, K. Kosanic, F. de Ory, F. J. Merino Fernandez, B. Gomez Rodriguez, [Zika virus infection acquired through sexual contact: first documented case of local transmission in Spain]. Emergencias 29, 290–291 (2017).

3. Q. Zhang et al., Spread of Zika virus in the Americas. Proc Natl Acad Sci U S A 114, E4334–E4343 (2017).

4. L. M. Araujo, M. L. Ferreira, O. J. Nascimento, Guillain-Barre syndrome associated with the Zika virus outbreak in Brazil. Arq Neuropsiquiatr 74, 253–255 (2016).

5. T. C. Pierson, B. S. Graham, Zika Virus: Immunity and Vaccine Development. Cell 167, 625–631 (2016).

6. D. Sirohi et al., The 3.8 A resolution cryo-EM structure of Zika virus. Science 352, 467–470 (2016).

7. A. Wang, S. Thurmond, L. Islas, K. Hui, R. Hai, Zika virus genome biology and molecular pathogenesis. Emerg Microbes Infect 6, e13 (2017).

8. X. Xu et al., Contribution of intertwined loop to membrane association revealed by Zika virus full-length NS1 structure. EMBO J 35, 2170–2178 (2016).

9. R. Hilgenfeld, Zika virus NS1, a pathogenicity factor with many faces. EMBO J 35, 2631–2633 (2016).

10. K. A. Dowd et al., Broadly Neutralizing Activity of Zika Virus-Immune Sera Identifies a Single Viral Serotype. Cell Rep 16, 1485–1491 (2016).

11. W. Sornjai, S. Ramphan, N. Wikan, P. Auewarakul, D. R. Smith, High correlation between Zika virus NS1 antibodies and neutralizing antibodies in selected serum samples from normal healthy Thais. Sci Rep 9, 13498 (2019).

12. L. Yu et al., Delineating antibody recognition against Zika virus during natural infection. JCI Insight 2, (2017).

13. M. J. Bailey et al., Antibodies Elicited by an NS1-Based Vaccine Protect Mice against Zika Virus. MBio 10, (2019).

14. T. Hampton, DNA Vaccine Protects Monkeys Against Zika Virus Infection. JAMA 316, 1755 (2016).

15. K. A. Dowd et al., Rapid development of a DNA vaccine for Zika virus. Science 354, 237–240 (2016).

16. M. S. Diamond, J. E. Ledgerwood, T. C. Pierson, Zika Virus Vaccine Development: Progress in the Face of New Challenges. Annu Rev Med 70, 121–135 (2019).

17. G. A. Poland, I. G. Ovsyannikova, R. B. Kennedy, Zika Vaccine Development: Current Status. Mayo Clin Proc 94, 2572–2586 (2019).

18. M. R. Gaudinski et al., Safety, tolerability, and immunogenicity of two Zika virus DNA vaccine candidates in healthy adults: randomised, open-label, phase 1 clinical trials. Lancet 391, 552–562 (2018).

19. P. Tebas et al., Safety and Immunogenicity of an Anti-Zika Virus DNA Vaccine - Preliminary Report. N Engl J Med, (2017).

20. K. Modjarrad et al., Preliminary aggregate safety and immunogenicity results from three trials of a purified inactivated Zika virus vaccine candidate: phase 1, randomised, double-blind, placebo-controlled clinical trials. Lancet 391, 563–571 (2018).

21. C. Lopez-Camacho et al., Assessment of Immunogenicity and Efficacy of a Zika Vaccine Using Modified Vaccinia Ankara Virus as Carriers. Pathogens 8, (2019).

22. P. Perez et al., A Vaccine Based on a Modified Vaccinia Virus Ankara Vector Expressing Zika Virus Structural Proteins Controls Zika Virus Replication in Mice. Sci Rep 8, 17385 (2018).

23. A. C. Brault et al., A Zika Vaccine Targeting NS1 Protein Protects Immunocompetent Adult Mice in a Lethal Challenge Model. Sci Rep 7, 14769 (2017).

24. J. M. Richner et al., Vaccine Mediated Protection Against Zika Virus-Induced Congenital Disease. Cell 170, 273–283 e212 (2017).

25. K. K. A. Van Rompay et al., DNA vaccination before conception protects Zika virus-exposed pregnant macaques against prolonged viremia and improves fetal outcomes. Sci Transl Med 11, (2019).

26. R. A. Larocca et al., Adenovirus Vector-Based Vaccines Confer Maternal-Fetal Protection against Zika Virus Challenge in Pregnant IFN-alphabetaR(-/-) Mice. Cell Host Microbe 26, 591–600 e594 (2019).

27. C. Verheust, M. Goossens, K. Pauwels, D. Breyer, Biosafety aspects of modified vaccinia virus Ankara (MVA)-based vectors used for gene therapy or vaccination. Vaccine 30, 2623–2632 (2012).

28. WHO:, in WHO Weekly Epidemiological Record of 12 July 2019. (2019).

29. A. Suy et al., Prolonged Zika Virus Viremia during Pregnancy. N Engl J Med 375, 2611–2613 (2016).

30. D. E. Griffin, Measles Vaccine. Viral Immunol 31, 86–95 (2018).

31. A. P. Fiebelkorn et al., Measles Virus Neutralizing Antibody Response, Cell-Mediated Immunity, and Immunoglobulin G Antibody Avidity Before and After Receipt of a Third Dose of Measles, Mumps, and Rubella Vaccine in Young Adults. J Infect Dis 213, 1115–1123 (2016).

32. S. A. Rasmussen, D. J. Jamieson, What Obstetric Health Care Providers Need to Know About Measles and Pregnancy. Obstet Gynecol 126, 163–170 (2015).

33. (2019).

34. R. R. Marty, M. C. Knuchel, T. N. Morin, H. Y. Naim, An immune competent mouse model for the characterization of recombinant measles vaccines. Hum Vaccin Immunother 11, 83–90 (2015).

35. M. Mura et al., hCD46 receptor is not required for measles vaccine Schwarz strain replication in vivo: Type-I IFN is the species barrier in mice. Virology 524, 151–159 (2018).

36. K. E. Stephenson et al., Safety and immunogenicity of a Zika purified inactivated virus vaccine given via standard, accelerated, or shortened schedules: a single-centre, double-blind, sequential-group, randomised, placebo-controlled, phase 1 trial. Lancet Infect Dis 20, 1061–1070 (2020).

37. J. L. Slon Campos et al., DNA-immunisation with dengue virus E protein domains I/II, but not domain III, enhances Zika, West Nile and Yellow Fever virus infection. PLoS One 12, e0181734 (2017).

38. A. P. S. Rathore, W. A. A. Saron, T. Lim, N. Jahan, A. L. St John, Maternal immunity and antibodies to dengue virus promote infection and Zika virus-induced microcephaly in fetuses. Sci Adv 5, eaav3208 (2019).

39. S. Gupta et al., The Neonatal Fc receptor (FcRn) enhances human immunodeficiency virus type 1 (HIV-1) transcytosis across epithelial cells. PLoS Pathog 9, e1003776 (2013).

40. M. G. Zimmerman et al., Cross-Reactive Dengue Virus Antibodies Augment Zika Virus Infection of Human Placental Macrophages. Cell Host Microbe 24, 731–742 e736 (2018).

41. A. Gordon et al., Prior dengue virus infection and risk of Zika: A pediatric cohort in Nicaragua. PLoS Med 16, e1002726 (2019).

42. P. Pantoja et al., Zika virus pathogenesis in rhesus macaques is unaffected by pre-existing immunity to dengue virus. Nat Commun 8, 15674 (2017).

43. S. Tripathi et al., A novel Zika virus mouse model reveals strain specific differences in virus pathogenesis and host inflammatory immune responses. PLoS Pathog 13, e1006258 (2017).

44. G. Young et al., Complete Protection in Macaques Conferred by Purified Inactivated Zika Vaccine: Defining a Correlate of Protection. Sci Rep 10, 3488 (2020).

45. A. O. Hassan et al., A Gorilla Adenovirus-Based Vaccine against Zika Virus Induces Durable Immunity and Confers Protection in Pregnancy. Cell Rep 28, 2634–2646 e2634 (2019).

46. C. Shan et al., A single-dose live-attenuated vaccine prevents Zika virus pregnancy transmission and testis damage. Nat Commun 8, 676 (2017).

47. J. L. Slon-Campos et al., A protective Zika virus E-dimer-based subunit vaccine engineered to abrogate antibody-dependent enhancement of dengue infection. Nat Immunol 20, 1291–1298 (2019).

48. M. J. Bailey et al., Human antibodies targeting Zika virus NS1 provide protection against disease in a mouse model. Nat Commun 9, 4560 (2018).

49. B. Grubor-Bauk et al., NS1 DNA vaccination protects against Zika infection through T cell-mediated immunity in immunocompetent mice. Sci Adv 5, eaax2388 (2019).

50. J. Emanuel et al., A VSV-based Zika virus vaccine protects mice from lethal challenge. Sci Rep 8, 11043 (2018).

51. A. Li et al., A Zika virus vaccine expressing premembrane-envelope-NS1 polyprotein. Nat Commun 9, 3067 (2018).

52. R. A. Larocca et al., Vaccine protection against Zika virus from Brazil. Nature 536, 474–478 (2016).

53. H. Garg, M. Sedano, G. Plata, E. B. Punke, A. Joshi, Development of Virus-Like-Particle Vaccine and Reporter Assay for Zika Virus. J Virol 91, (2017).

54. K. Xu et al., Recombinant Chimpanzee Adenovirus Vaccine AdC7-M/E Protects against Zika Virus Infection and Testis Damage. J Virol 92, (2018).

55. B. W. Jagger et al., Protective Efficacy of Nucleic Acid Vaccines Against Transmission of Zika Virus During Pregnancy in Mice. J Infect Dis 220, 1577–1588 (2019).

56. J. R. Keeffe et al., A Combination of Two Human Monoclonal Antibodies Prevents Zika Virus Escape Mutations in Non-human Primates. Cell Rep 25, 1385–1394 e1387 (2018).

57. C. Nurnberger, B. S. Bodmer, A. H. Fiedler, G. Gabriel, M. D. Muhlebach, A Measles Virus-Based Vaccine Candidate Mediates Protection against Zika Virus in an Allogeneic Mouse Pregnancy Model. J Virol 93, (2019).

58. C. Shan et al., Maternal vaccination and protective immunity against Zika virus vertical transmission. Nat Commun 10, 5677 (2019).

59. D. F. Robbiani et al., Risk of Zika microcephaly correlates with features of maternal antibodies. J Exp Med 216, 2302–2315 (2019).

60. D. F. Robbiani et al., Recurrent Potent Human Neutralizing Antibodies to Zika Virus in Brazil and Mexico. Cell 169, 597–609 e511 (2017).

61. M. Hassert, M. G. Harris, J. D. Brien, A. K. Pinto, Identification of Protective CD8 T Cell Responses in a Mouse Model of Zika Virus Infection. Front Immunol 10, 1678 (2019).

62. M. Hassert et al., CD4+T cells mediate protection against Zika associated severe disease in a mouse model of infection. PLoS Pathog 14, e1007237 (2018).

63. I. G. Ovsyannikova, N. Dhiman, R. M. Jacobson, R. A. Vierkant, G. A. Poland, Frequency of measles virus-specific CD4+ and CD8+ T cells in subjects seronegative or highly seropositive for measles vaccine. Clin Diagn Lab Immunol 10, 411–416 (2003).

64. C. Shan et al., An Infectious cDNA Clone of Zika Virus to Study Viral Virulence, Mosquito Transmission, and Antiviral Inhibitors. Cell Host Microbe 19, 891–900 (2016).

65. H. Zhao et al., Structural Basis of Zika Virus-Specific Antibody Protection. Cell 166, 1016–1027 (2016).

66. M. S. Lee, B. Cohen, J. Hand, D. J. Nokes, A simplified and standardized neutralization enzyme immunoassay for the quantification of measles neutralizing antibody. J Virol Methods 78, 209–217 (1999).

67. H. B. Shah, K. A. Koelsch, B-Cell ELISPOT: For the Identification of Antigen-Specific Antibody-Secreting Cells. Methods Mol Biol 1312, 419–426 (2015).

