## supplement for "Measles-based Zika vaccine induces long-term immunity and requires NS1 antibodies to protect the female reproductive tract in the hCD46 IFNα/β receptor knockout mice" (from another version of this preprint: version 1, 2020-09-17)

```
MluI BsiWI
5' ACGCGTGGAATCCTCCCGTACGGCCACCATGGGGGCTGATACAAGCATTGGCATCATTGGGCTGCTGCTGACCACAGCTATGGCAGCTGAGGTGACAAGGCGCGGCTCCGCCTACTATATGTACCTGGACCGGAATGATGCTGGCGAAGCAATCAGCTTCCCAACTACAC 170
3' TGCGCACCTTAGGAGGGCATGCCGGTGGTACCCCCGACTATGTTCGTAACCGTAGTAACCCGACGACGACTGGTGTCGATACCGTCGACTCCACTGTTCCGCGCCGAGGCGGATGATATACATGGACCTGGCCTTACTACGACCGCTTCGTTAGTCGAAGGGTTGATGTG
Signal Peptide | precursor protein
M G A D T S I G I I G L L L T T A M A A E V T R R G S A Y Y M Y L D R N D A G E A I S F P T T
1 2 3 4 5 6 7 8 9 10 11 12 13 14 15 16 17 18 19 20 21 22 23 24 25 26 27 28 29 30 31 32 33 34 35 36 37 38 39 40 41 42 43 44 45 46 47

5' TGGGCATGAACAAGTGCTACATCCAGATTATGGACCTGGGCCACATGTGCGATGCCACCATGTCCTATGAGTGTCCAATGCTGGACGAGGGGGTGGAACCCGACGATGTCGATTGCTGGTGTAATACAACTAGTACATGGGTGGTCTACGGCACTTGTCACCATAAGAAA 340
3' ACCCGTACTTGTTCACGATGTAGGTCTAATACCTGGACCCGGTGTACACGCTACGGTGGTACAGGATACTCACAGGTTACGACCTGCTCCCCCACCTTGGGCTGCTACAGCTAACGACCACATTATGTTGATCATGTACCCACCAGATGCCGTGAACAGTGGTATTCTTT
precursor protein
L G M N K C Y I Q I M D L G H M C D A T M S Y E C P M L D E G V E P D D V D C W C N T T S T W V V Y G T C H H K K
48 49 50 51 52 53 54 55 56 57 58 59 60 61 62 63 64 65 66 67 68 69 70 71 72 73 74 75 76 77 78 79 80 81 82 83 84 85 86 87 88 89 90 91 92 93 94 95 96 97 98 99 100 101 102 103 104

5' GGGGAAGCTCGACGGTCAAGAAGGGCAGTGACCCTGCCCTCACACAGCACAAGGAAGCTCCAGACCCGCTCTCAGACATGGCTGGAGAGTAGAGAATATACAAAACATCTGATCAGAGTGGAGAACTGGATCTTCCGGAATCCAGGATTCGCACTGGCAGCCGCTGCAAT 510
3' CCCCTTCGAGCTGCCAGTTCTTCCCGTCACTGGGACGGGAGTGTGTCGTGTTCCTTCGAGGTCTGGGCGAGAGTCTGTACCGACCTCTCATCTCTTATATGTTTTGTAGACTAGTCTCACCTCTTGACCTAGAAGGCCTTAGGTCCTAAGCGTGACCGTCGGCGACGTTA
precursor protein | Membrane protein
G E A R R S R R A V T L P S H S T R K L Q T R S Q T W L E S R E Y T K H L I R V E N W I F R N P G F A L A A A A I
105 106 107 108 109 110 111 112 113 114 115 116 117 118 119 120 121 122 123 124 125 126 127 128 129 130 131 132 133 134 135 136 137 138 139 140 141 142 143 144 145 146 147 148 149 150 151 152 153 154 155 156 157 158 159 160 161

5' CGCATGGCTGCTGGGCAGCTCCACCTCTCAGAAAGTGATCTACCTGGTCATGATCCTGCTGATTGCCCCCGCTTATTCCATCCGCTGCATTGGGGTGTCTAATCGAGACTTCGTCGAGGGAATGAGTGGCGGGACCTGGGTGGATGTGGTCCTGGAACACGGAGGCTGCG 680
3' GCGTACCGACGACCCGTCGAGGTGGAGAGTCTTTCACTAGATGGACCAGTACTAGGACGACTAACGGGGGCGAATAAGGTAGGCGACGTAACCCCACAGATTAGCTCTGAAGCAGCTCCCTTACTCACCGCCCTGGACCCACCTACACCAGGACCTTGTGCCTCCGACGC
Membrane protein | Envelope
A W L L G S S T S Q K V I Y L V M I L L I A P A Y S I R C I G V S N R D F V E G M S G G T W V D V V L E H G G C
162 163 164 165 166 167 168 169 170 171 172 173 174 175 176 177 178 179 180 181 182 183 184 185 186 187 188 189 190 191 192 193 194 195 196 197 198 199 200 201 202 203 204 205 206 207 208 209 210 211 212 213 214 215 216 217

5' TGACAGTGATGGCTCAGGACAAGCCTACTGTGGACATCGAGCTGGTGACCACAACTGTCAGTAACATGGCCGAGGTGAGGTCATACTGCTATGAAGCCTCCATTTCTGACATGGCTAGTGATTCACGCTGTCCAACCCAGGGCGAGGCCTACCTGGACAAGCAGAGCGAT 850
3' ACTGTCACTACCGAGTCCTGTTCGGATGACACCTGTAGCTCGACCACTGGTGTTGACAGTCATTGTACCGGCTCCACTCCAGTATGACGATACTTCGGAGGTAAAGACTGTACCGATCACTAAGTGCGACAGGTTGGGTCCCGCTCCGGATGGACCTGTTCGTCTCGCTA
Envelope
V T V M A Q D K P T V D I E L V T T T V S N M A E V R S Y C Y E A S I S D M A S D S R C P T Q G E A Y L D K Q S D
218 219 220 221 222 223 224 225 226 227 228 229 230 231 232 233 234 235 236 237 238 239 240 241 242 243 244 245 246 247 248 249 250 251 252 253 254 255 256 257 258 259 260 261 262 263 264 265 266 267 268 269 270 271 272 273 274

5' ACTCAGTACGTGTGCAAACGAACCCTGGTGGACCGGGGCTGGGGGAATGGATGTGGCCTGTTTGGGAAGGGCAGCCTGGTGACCTGCGCCAAATTCGCTTGTAGCAAGAAAATGACCGGCAAGTCCATCCAGCCTGAGAACCTGGAATACAGGATTATGCTGTCTGTGCA 1020
3' TGAGTCATGCACACGTTTGCTTGGGACCACCTGGCCCCGACCCCCTTACCTACACCGGACAAACCCTTCCCGTCGGACCACTGGACGCGGTTTAAGCGAACATCGTTCTTTTACTGGCCGTTCAGGTAGGTCGGACTCTTGGACCTTATGTCCTAATACGACAGACACGT
Envelope
T Q Y V C K R T L V D R G W G N G C G L F G K G S L V T C A K F A C S K K M T G K S I Q P E N L E Y R I M L S V H
275 276 277 278 279 280 281 282 283 284 285 286 287 288 289 290 291 292 293 294 295 296 297 298 299 300 301 302 303 304 305 306 307 308 309 310 311 312 313 314 315 316 317 318 319 320 321 322 323 324 325 326 327 328 329 330 331

5' CGGCAGCCAGCATTCTGGCATGATCGTCAACGACACTGGCCACGAGACAGATGAAAATCGAGCCAAAGTGGAGATTACTCCTAACTCCCCAAGAGCAGAAGCCACCCTGGGGGGATTTGGATCTCTGGGCCTGGACTGCGAGCCACGAACTGGCCTGGACTTCTCCGATC 1190
3' GCCGTCGGTCGTAAGACCGTACTAGCAGTTGCTGTGACCGGTGCTCTGTCTACTTTTAGCTCGGTTTCACCTCTAATGAGGATTGAGGGGTTCTCGTCTTCGGTGGGACCCCCCTAAACCTAGAGACCCGGACCTGACGCTCGGTGCTTGACCGGACCTGAAGAGGCTAG
Envelope
G S Q H S G M I V N D T G H E T D E N R A K V E I T P N S P R A E A T L G G F G S L G L D C E P R T G L D F S D
332 333 334 335 336 337 338 339 340 341 342 343 344 345 346 347 348 349 350 351 352 353 354 355 356 357 358 359 360 361 362 363 364 365 366 367 368 369 370 371 372 373 374 375 376 377 378 379 380 381 382 383 384 385 386 387

5' TGTACTATCTGACCATGAACAATAAGCACTGGCTGGTGCATAAAGAATGGTTTCACGACATCCCACTGCCCTGGCACGCCGGAGCAGATACTGGCACCCCTCACTGGAACAATAAGGAGGCTCTGGTGGAGTTCAAGGATGCCCATGCTAAACGACAGACCGTGGTCGTG 1360
3' ACATGATAGACTGGTACTTGTTATTCGTGACCGACCACGTATTTCTTACCAAAGTGCTGTAGGGTGACGGGACCGTGCGGCCTCGTCTATGACCGTGGGGAGTGACCTTGTTATTCCTCCGAGACCACCTCAAGTTCCTACGGGTACGATTTGCTGTCTGGCACCAGCAC
Envelope
L Y Y L T M N N K H W L V H K E W F H D I P L P W H A G A D T G T P H W N N K E A L V E F K D A H A K R Q T V V V
388 389 390 391 392 393 394 395 396 397 398 399 400 401 402 403 404 405 406 407 408 409 410 411 412 413 414 415 416 417 418 419 420 421 422 423 424 425 426 427 428 429 430 431 432 433 434 435 436 437 438 439 440 441 442 443 444

5' CTGGGGAGCCAGGAGGGAGCAGTGCACACAGCACTGGCCGGCGCTCTGGAGGCAGAAATGGACGGGGCCAAGGGAAGACTGTCTAGTGGGCATCTGAAATGCCGGCTGAAGATGGATAAACTGAGACTGAAGGGAGTGAGCTACTCCCTGTGCACCGCCGCCTTCACCTT 1530
3' GACCCCTCGGTCCTCCCTCGTCACGTGTGTCGTGACCGGCCGCGAGACCTCCGTCTTTACCTGCCCCGGTTCCCTTCTGACAGATCACCCGTAGACTTTACGGCCGACTTCTACCTATTTGACTCTGACTTCCCTCACTCGATGAGGGACACGTGGCGGCGGAAGTGGAA
Envelope
L G S Q E G A V H T A L A G A L E A E M D G A K G R L S S G H L K C R L K M D K L R L K G V S Y S L C T A A F T F
445 446 447 448 449 450 451 452 453 454 455 456 457 458 459 460 461 462 463 464 465 466 467 468 469 470 471 472 473 474 475 476 477 478 479 480 481 482 483 484 485 486 487 488 489 490 491 492 493 494 495 496 497 498 499 500 501

5' CACCAAAATCCCAGCCGAGACTCTGCACGGCACCGTCACAGTGGAAGTCCAGTATGCTGGCACTGACGGCCCTTGTAAGGTGCCTGCACAGATGGCCGTCGATATGCAGACTCTGACCCCAGTGGGCCGGCTGATCACCGCAAATCCTGTCATTACAGAGAGCACTGAAA 1700
3' GTGGTTTTAGGGTCGGCTCTGAGACGTGCCGTGGCAGTGTCACCTTCAGGTCATACGACCGTGACTGCCGGGAACATTCCACGGACGTGTCTACCGGCAGCTATACGTCTGAGACTGGGGTCACCCGGCCGACTAGTGGCGTTTAGGACAGTAATGTCTCTCGTGACTTT
Envelope
T K I P A E T L H G T V T V E V Q Y A G T D G P C K V P A Q M A V D M Q T L T P V G R L I T A N P V I T E S T E
502 503 504 505 506 507 508 509 510 511 512 513 514 515 516 517 518 519 520 521 522 523 524 525 526 527 528 529 530 531 532 533 534 535 536 537 538 539 540 541 542 543 544 545 546 547 548 549 550 551 552 553 554 555 556 557

5' ACTCCAAAATGATGCTGGAGCTGGACCCCCCTTTTGGGGATAGCTATATCGTGATTGGCGTCGGGGAAAAGAAAATCACACACCATTGGCACCGGAGCGGCAGTACCATTGGGAAGGCTTTTGAGGCAACAGTGCGCGGCGCCAAACGAATGGCTGTCCTGGGAGACACC 1870
3' TGAGGTTTTACTACGACCTCGACCTGGGGGGAAAACCCCTATCGATATAGCACTAACCGCAGCCCCTTTTCTTTTAGTGTGTGGTAACCGTGGCCTCGCCGTCATGGTAACCCTTCCGAAAACTCCGTTGTCACGCGCCGCGGTTTGCTTACCGACAGGACCCTCTGTGG
Envelope
N S K M M L E L D P P F G D S Y I V I G V G E K K I T H H W H R S G S T I G K A F E A T V R G A K R M A V L G D T
558 559 560 561 562 563 564 565 566 567 568 569 570 571 572 573 574 575 576 577 578 579 580 581 582 583 584 585 586 587 588 589 590 591 592 593 594 595 596 597 598 599 600 601 602 603 604 605 606 607 608 609 610 611 612 613 614

5' GCTTGGGATTTCGGCTCTGTGGGCGGGGCATTCAACAGTCTGGGAAAGGGCATCCATCAGATTTTCGGAGCAGCCTTTAAATCCCTGTTCGGAGGCATGAGTTGGTTCTCACAGATCCTGATTGGCACACTGCTGATGTGGCTGGGGCTGAACACTAAGAATGGATCAAT 2040
3' CGAACCCTAAAGCCGAGACACCCGCCCCGTAAGTTGTCAGACCCTTTCCCGTAGGTAGTCTAAAAGCCTCGTCGGAAATTTAGGGACAAGCCTCCGTACTCAACCAAGAGTGTCTAGGACTAACCGTGTGACGACTACACCGACCCCGACTTGTGATTCTTACCTAGTTA
Envelope
TM1 | TM2
A W D F G S V G G A F N S L G K G I H Q I F G A A F K S L F G G M S W F S Q I L I G T L L M W L G L N T K N G S I
615 616 617 618 619 620 621 622 623 624 625 626 627 628 629 630 631 632 633 634 635 636 637 638 639 640 641 642 643 644 645 646 647 648 649 650 651 652 653 654 655 656 657 658 659 660 661 662 663 664 665 666 667 668 669 670 671

AatII
5' CAGCCTGATGTGCCTGGCTCTGGGGGGGGTCCTGATTTTTCTGTCCACCGCAGTGTCCGCATAAATGACGTC 2112
3' GTCGGACTACACGGACCGAGACCCCCCCCAGGACTAAAAAGACAGGTGGCGTCACAGGCGTATTTACTGCAG
Envelope
TM2
S L M C L A L G G V L I F L S T A V S A .
672 673 674 675 676 677 678 679 680 681 682 683 684 685 686 687 688 689 690 691 692
```

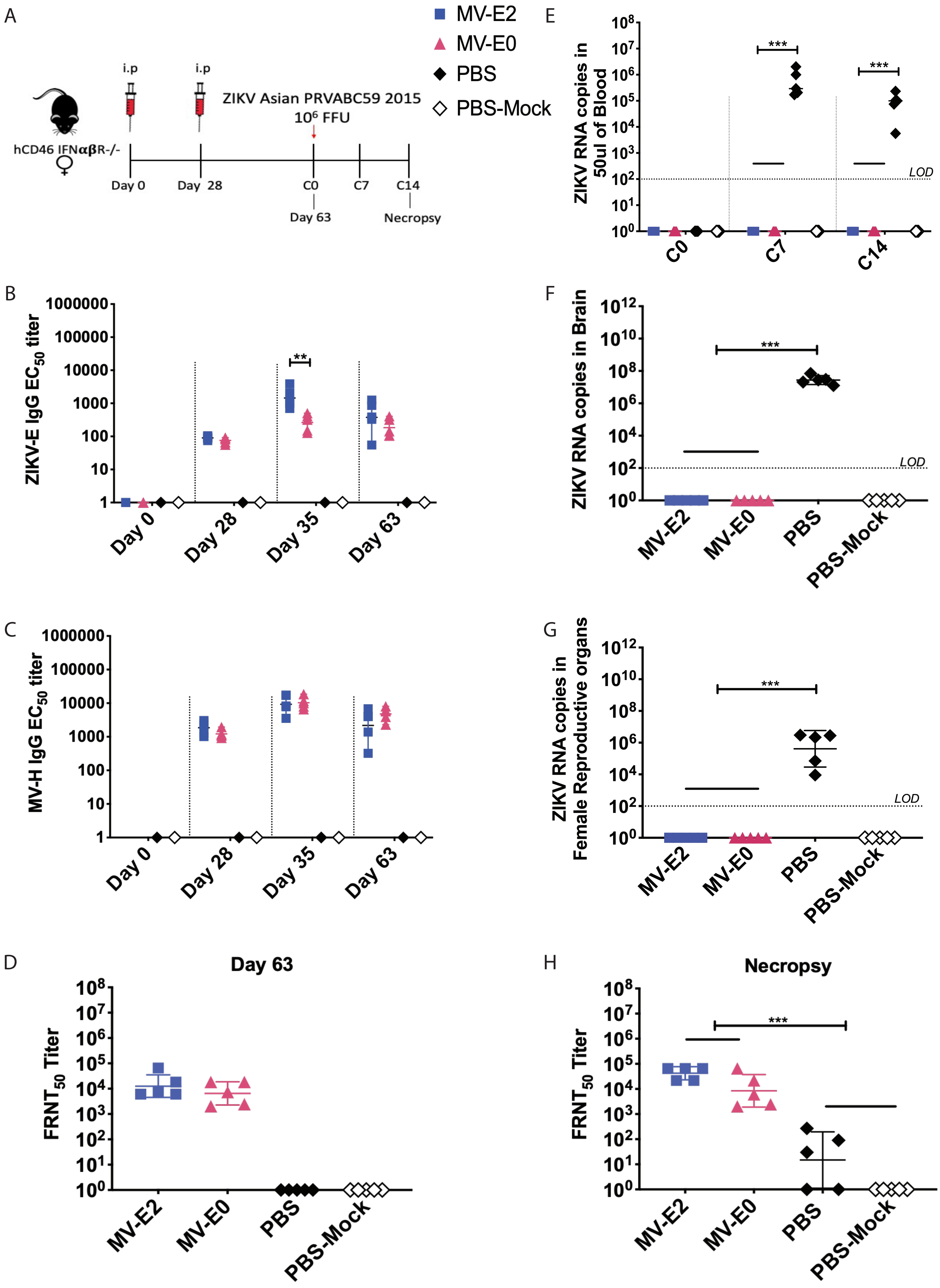
A
i.p
i.p
ZIKV Asian PRVABC59 2015
10^6 FFU
hCD46 IFNαβR-/-
♀
Day 0
Day 28
C0
C7
C14
Day 63
Necropsy
MV-E2
MV-E0
PBS
PBS-Mock
B
ZIKV-E IgG EC50 titer
1000000
100000
10000
1000
100
10
1
**
Day 0
Day 28
Day 35
Day 63
C
MV-H IgG EC50 titer
1000000
100000
10000
1000
100
10
1
Day 0
Day 28
Day 35
Day 63
D
Day 63
FRNT50 Titer
10^8
10^7
10^6
10^5
10^4
10^3
10^2
10^1
10^0
MV-E2
MV-E0
PBS
PBS-Mock
E
ZIKV RNA copies in 50ul of Blood
10^8
10^7
10^6
10^5
10^4
10^3
10^2
10^1
10^0
***
***
LOD
C0
C7
C14
F
ZIKV RNA copies in Brain
10^12
10^10
10^8
10^6
10^4
10^2
10^0
***
LOD
MV-E2
MV-E0
PBS
PBS-Mock
G
ZIKV RNA copies in Female Reproductive organs
10^12
10^10
10^8
10^6
10^4
10^2
10^0
***
LOD
MV-E2
MV-E0
PBS
PBS-Mock
H
Necropsy
FRNT50 Titer
10^8
10^7
10^6
10^5
10^4
10^3
10^2
10^1
10^0
***
MV-E2
MV-E0
PBS
PBS-Mock

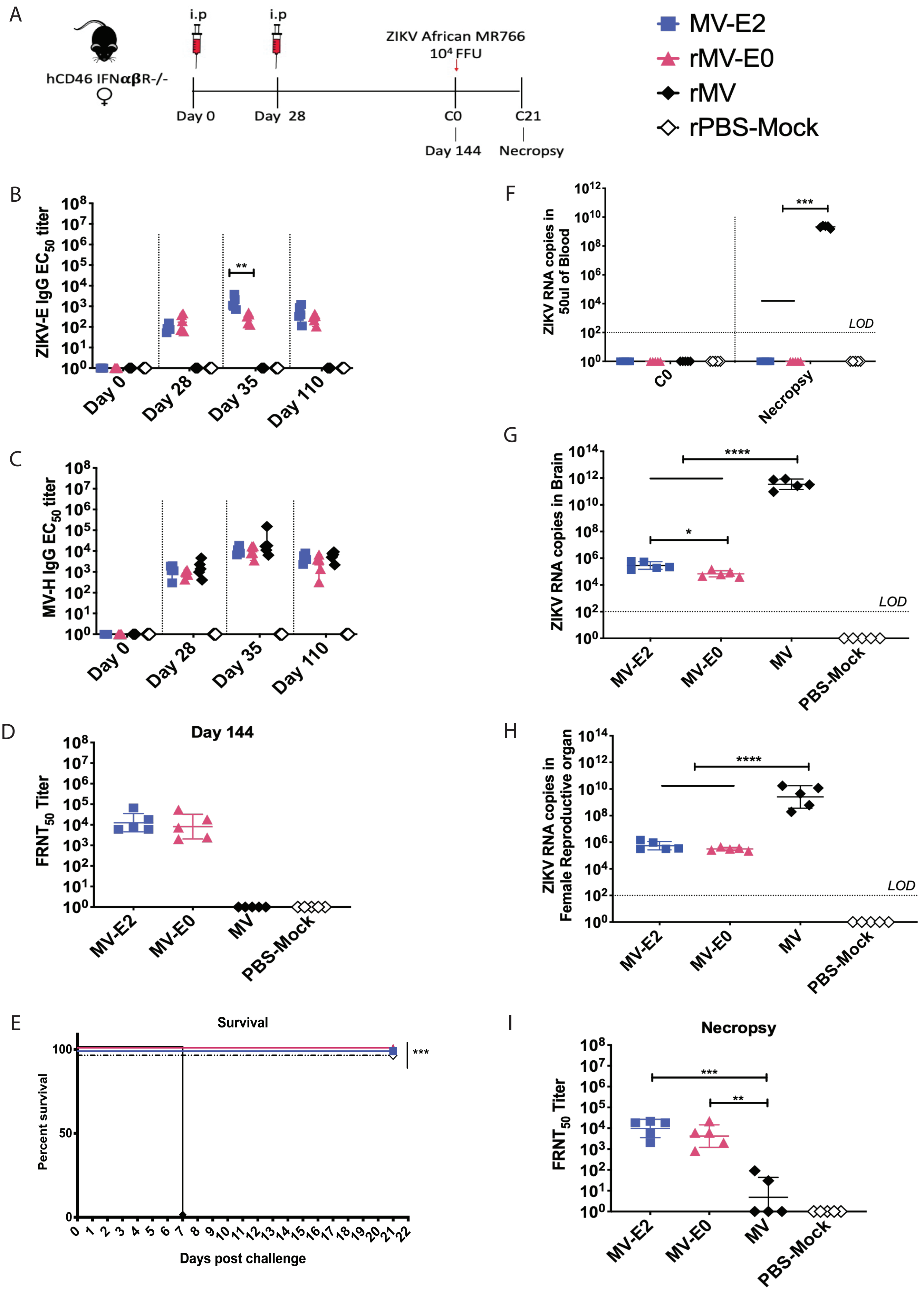

A
i.p
i.p
hCD46 IFNαβR-/-
ZIKV African MR766
10⁴ FFU
Day 0
Day 28
C0
C21
Day 144
Necropsy
MV-E2
rMV-E0
rMV
rPBS-Mock
B
ZIKV-E IgG EC50 titer
Day 0
Day 28
Day 35
Day 110
C
MV-H IgG EC50 titer
D
Day 144
FRNT50 Titer
MV-E2
MV-E0
MV
PBS-Mock
E
Survival
Percent survival
Days post challenge
F
ZIKV RNA copies in 50ul of Blood
LOD
C0
Necropsy
G
ZIKV RNA copies in Brain
H
ZIKV RNA copies in Female Reproductive organ
I
Necropsy

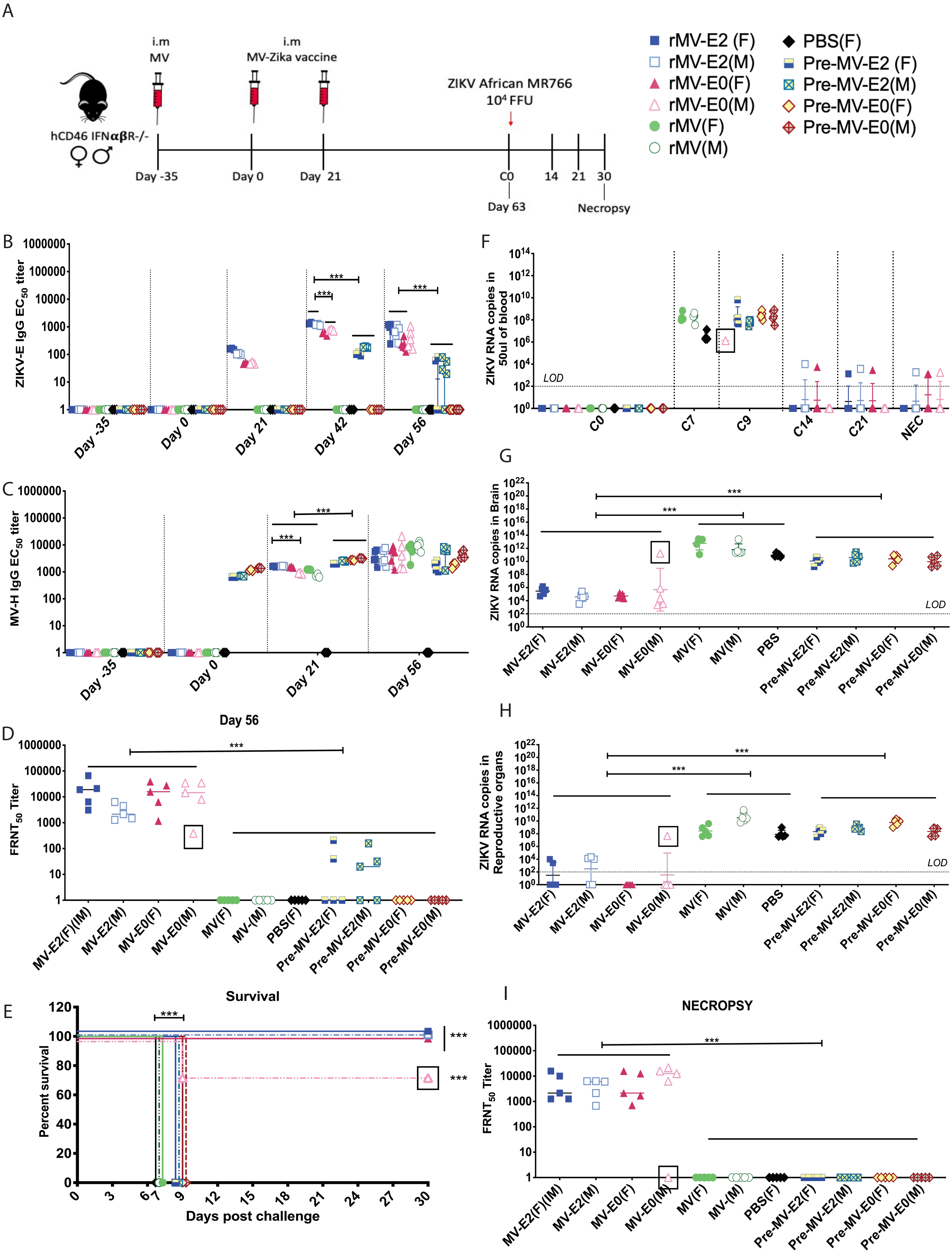

A
i.m
MV
i.m
MV-Zika vaccine
ZIKV African MR766
10⁴ FFU
hCD46 IFNαβR-/-
♀ ♂
Day -35
Day 0
Day 21
C0
14
21
30
Day 63
Necropsy
rMV-E2 (F)
rMV-E2(M)
rMV-E0(F)
rMV-E0(M)
rMV(F)
rMV(M)
PBS(F)
Pre-MV-E2 (F)
Pre-MV-E2(M)
Pre-MV-E0(F)
Pre-MV-E0(M)
B
ZIKV-E IgG EC50 titer
Day -35
Day 0
Day 21
Day 42
Day 56
***
C
MV-H IgG EC50 titer
Day -35
Day 0
Day 21
Day 56
***
D
Day 56
FRNT50 Titer
MV-E2(F)(IM)
MV-E2(M)
MV-E0(F)
MV-E0(M)
MV(F)
MV-(M)
PBS(F)
Pre-MV-E2(F)
Pre-MV-E2(M)
Pre-MV-E0(F)
Pre-MV-E0(M)
***
E
Survival
Percent survival
Days post challenge
***
F
ZIKV RNA copies in 50ul of blood
LOD
C0
C7
C9
C14
C21
NEC
G
ZIKV RNA copies in Brain
LOD
MV-E2(F)
MV-E2(M)
MV-E0(F)
MV-E0(M)
MV(F)
MV(M)
PBS
Pre-MV-E2(F)
Pre-MV-E2(M)
Pre-MV-E0(F)
Pre-MV-E0(M)
***
H
ZIKV RNA copies in Reproductive organs
LOD
***
I
NECROPSY
FRNT50 Titer
***

```
                                       MluI
                                         BsiWI
5' AGGAACCAGGTCCACACAGAGTGATACGCGTACGGGCCACCATGGGGGCTGATACAAGCGTCGGAATCGTGGGGCTGCTGCTGACAACCGCAATGGCTGCTGAAGTCACAAGAAGAGGGTCTGCCTACTACATGTATCTGGACAGGAATG 150
3' TCCTTGGTCCAGGTGTGTCTCACTATGCGCATGCCCGGTGGTACCCCCGACTATGTTCGCAGCCTTAGCACCCCGACGACGACTGTTGGCGTTACCGACGACTTCAGTGTTCTTCTCCCAGACGGATGATGTACATAGACCTGTCCTTAC
Signal Peptide | Precursor protein
M G A D T S V G I V G L L L T T A M A A E V T R R G S A Y Y M Y L D R N
1 2 3 4 5 6 7 8 9 10 11 12 13 14 15 16 17 18 19 20 21 22 23 24 25 26 27 28 29 30 31 32 33 34 35 36

5' ATGCCGGCGAGGCCATCTCCTTTCCTACCACACTGGGCATGAACAAGTGCTACATCCAGATCATGGACCTGGGCCACATGTGCGATGCCACCATGTCTTATGAGTGTCCCATGCTGGACGAGGGCGTGGAGCCTGACGATGTGGATTGCT 300
3' TACGGCCGCTCCGGTAGAGGAAAGGATGGTGTGACCCGTACTTGTTCACGATGTAGGTCTAGTACCTGGACCCGGTGTACACGCTACGGTGGTACAGAATACTCACAGGGTACGACCTGCTCCCGCACCTCGGACTGCTACACCTAACGA
Precursor protein
D A G E A I S F P T T L G M N K C Y I Q I M D L G H M C D A T M S Y E C P M L D E G V E P D D V D C
37 38 39 40 41 42 43 44 45 46 47 48 49 50 51 52 53 54 55 56 57 58 59 60 61 62 63 64 65 66 67 68 69 70 71 72 73 74 75 76 77 78 79 80 81 82 83 84 85 86

5' GGTGTAACACCACAAGCACATGGGTGGTGTACGGCACCTGTCACCACAAGAAGGGAGAGGCCCGGCGGAGCCGGCGGGCCGTGACACTGCCAAGCCACTCCACCCGGAAGCTGCAGACAAGAAGCCAGACCTGGCTGGAGTCCCGCGAGT 450
3' CCACATTGTGGTGTTCGTGTACCCACCACATGCCGTGGACAGTGGTGTTCTTCCCTCTCCGGGCCGCCTCGGCCGCCCGGCACTGTGACGGTTCGGTGAGGTGGGCCTTCGACGTCTGTTCTTCGGTCTGGACCGACCTCAGGGCGCTCA
Precursor protein | Membrane Protein
W C N T T S T W V V Y G T C H H K K G E A R R S R R A V T L P S H S T R K L Q T R S Q T W L E S R E
87 88 89 90 91 92 93 94 95 96 97 98 99 100 101 102 103 104 105 106 107 108 109 110 111 112 113 114 115 116 117 118 119 120 121 122 123 124 125 126 127 128 129 130 131 132 133 134 135 136

5' ATACAAAGCACCTGATCAGGGTGGAGAACTGGATCTTTCGCAATCCAGGATTCGCACTGGCAGCAGCAGCAATCGCCTGGCTGCTGGGCAGCTCCACCTCTCAGAAAGTGATCTACCTGGTCATGATCCTGCTGATCGCCCCTGCCTATT 600
3' TATGTTTCGTGGACTAGTCCCACCTCTTGACCTAGAAAGCGTTAGGTCCTAAGCGTGACCGTCGTCGTCGTTAGCGGACCGACGACCCGTCGAGGTGGAGAGTCTTTCACTAGATGGACCAGTACTAGGACGACTAGCGGGGACGGATAA
Membrane Protein
Y T K H L I R V E N W I F R N P G F A L A A A A I A W L L G S S T S Q K V I Y L V M I L L I A P A Y
137 138 139 140 141 142 143 144 145 146 147 148 149 150 151 152 153 154 155 156 157 158 159 160 161 162 163 164 165 166 167 168 169 170 171 172 173 174 175 176 177 178 179 180 181 182 183 184 185 186

5' CCATCCGGTGCATCGGCGTGTCTAATAGAGACTTCGTGGAGGGAATGTCCGGAGGAACCTGGGTGGATGTGGTGCTGGAGCACGGCGGCTGCGTGACAGTGATGGCCCAGGACAAGCCTACCGTGGATATCGAGCTGGTGACCACAACCG 750
3' GGTAGGCCACGTAGCCGCACAGATTATCTCTGAAGCACCTCCCTTACAGGCCTCCTTGGACCCACCTACACCACGACCTCGTGCCGCCGACGCACTGTCACTACCGGGTCCTGTTCGGATGGCACCTATAGCTCGACCACTGGTGTTGGC
Envelope protein
S I R C I G V S N R D F V E G M S G G T W V D V V L E H G G C V T V M A Q D K P T V D I E L V T T T
187 188 189 190 191 192 193 194 195 196 197 198 199 200 201 202 203 204 205 206 207 208 209 210 211 212 213 214 215 216 217 218 219 220 221 222 223 224 225 226 227 228 229 230 231 232 233 234 235 236

5' TGAGCAACATGGCCGAGGTGAGGTCCTACTGCTATGAGGCCTCTATCAGCGACATGGCCTCCGATTCTCGCTGTCCAACCCAGGGCGAGGCCTACCTGGACAAGCAGAGCGATACACAGTACGTGTGCAAGAGGACCCTGGTGGACAGGG 900
3' ACTCGTTGTACCGGCTCCACTCCAGGATGACGATACTCCGGAGATAGTCGCTGTACCGGAGGCTAAGAGCGACAGGTTGGGTCCCGCTCCGGATGGACCTGTTCGTCTCGCTATGTGTCATGCACACGTTCTCCTGGGACCACCTGTCCC
Envelope protein
V S N M A E V R S Y C Y E A S I S D M A S D S R C P T Q G E A Y L D K Q S D T Q Y V C K R T L V D R
237 238 239 240 241 242 243 244 245 246 247 248 249 250 251 252 253 254 255 256 257 258 259 260 261 262 263 264 265 266 267 268 269 270 271 272 273 274 275 276 277 278 279 280 281 282 283 284 285 286

5' GATGGGGAAATGGATGTGGCCTGTTTGGCAAGGGCTCCCTGGTGACATGCGCCAAGTTCGCCTGTTCTAAGAAGATGACCGGCAAGAGCATCCAGCCCGAGAACCTGGAGTACCGGATCATGCTGAGCGTGCACGGCAGCCAGCACTCCG 1050
3' CTACCCCTTTACCTACACCGGACAAACCGTTCCCGAGGGACCACTGTACGCGGTTCAAGCGGACAAGATTCTTCTACTGGCCGTTCTCGTAGGTCGGGCTCTTGGACCTCATGGCCTAGTACGACTCGCACGTGCCGTCGGTCGTGAGGC
Envelope protein
G W G N G C G L F G K G S L V T C A K F A C S K K M T G K S I Q P E N L E Y R I M L S V H G S Q H S
287 288 289 290 291 292 293 294 295 296 297 298 299 300 301 302 303 304 305 306 307 308 309 310 311 312 313 314 315 316 317 318 319 320 321 322 323 324 325 326 327 328 329 330 331 332 333 334 335 336

5' GCATGATCGTGAACGACACAGGCCACGAGACAGATGAGAATCGGGCCAAGGTGGAGATCACACCAAACTCCCCAAGAGCAGAGGCCACCCTGGGAGGATTTGGCTCTCTGGGCCTGGACTGCGAGCCCAGAACAGGCCTGGACTTCTCCG 1200
3' CGTACTAGCACTTGCTGTGTCCGGTGCTCTGTCTACTCTTAGCCCGGTTCCACCTCTAGTGTGGTTTGAGGGGTTCTCGTCTCCGGTGGGACCCTCCTAAACCGAGAGACCCGGACCTGACGCTCGGGTCTTGTCCGGACCTGAAGAGGC
Envelope protein
G M I V N D T G H E T D E N R A K V E I T P N S P R A E A T L G G F G S L G L D C E P R T G L D F S
337 338 339 340 341 342 343 344 345 346 347 348 349 350 351 352 353 354 355 356 357 358 359 360 361 362 363 364 365 366 367 368 369 370 371 372 373 374 375 376 377 378 379 380 381 382 383 384 385 386

5' ATCTGTACTATCTGACCATGAACAATAAGCACTGGCTGGTGCACAAGGAGTGGTTTCACGACATCCCTCTGCCATGGCACGCAGGAGCAGATACAGGAACCCCTCACTGGAACAATAAGGAGGCCCTGGTGGAGTTCAAGGATGCCCACG 1350
3' TAGACATGATAGACTGGTACTTGTTATTCGTGACCGACCACGTGTTCCTCACCAAAGTGCTGTAGGGAGACGGTACCGTGCGTCCTCGTCTATGTCCTTGGGGAGTGACCTTGTTATTCCTCCGGGACCACCTCAAGTTCCTACGGGTGC
Envelope protein
D L Y Y L T M N N K H W L V H K E W F H D I P L P W H A G A D T G T P H W N N K E A L V E F K D A H
387 388 389 390 391 392 393 394 395 396 397 398 399 400 401 402 403 404 405 406 407 408 409 410 411 412 413 414 415 416 417 418 419 420 421 422 423 424 425 426 427 428 429 430 431 432 433 434 435 436

5' CCAAGAGGCAGACAGTGGTGGTGCTGGGCAGCCAGGAGGGAGCAGTGCACACCGCCCTGGCAGGCGCCCTGGAGGCCGAGATGGACGGAGCAAAGGGCCGCCTGTCTAGCGGCCACCTGAAGTGCAGGCTGAAGATGGATAAGCTGCGCC 1500
3' GGTTCTCCGTCTGTCACCACCACGACCCGTCGGTCCTCCCTCGTCACGTGTGGCGGGACCGTCCGCGGGACCTCCGGCTCTACCTGCCTCGTTTCCCGGCGGACAGATCGCCGGTGGACTTCACGTCCGACTTCTACCTATTCGACGCGG
Envelope protein
A K R Q T V V V L G S Q E G A V H T A L A G A L E A E M D G A K G R L S S G H L K C R L K M D K L R
437 438 439 440 441 442 443 444 445 446 447 448 449 450 451 452 453 454 455 456 457 458 459 460 461 462 463 464 465 466 467 468 469 470 471 472 473 474 475 476 477 478 479 480 481 482 483 484 485 486

5' TGAAGGGCGTGAGCTACTCCCTGTGCACAGCCGCCTTCACCTTCACCAAGATCCCTGCCGAGACACTGCACGGCACAGTGACCGTGGAGCTGCAGTATGCAGGAACAGACGGACCATGCAAGGTGCCTGCACAGATGGCCGTGGATATGC 1650
3' ACTTCCCGCACTCGATGAGGGACACGTGTCGGCGGAAGTGGAAGTGGTTCTAGGGACGGCTCTGTGACGTGCCGTGTCACTGGCACCTCGACGTCATACGTCCTTGTCTGCCTGGTACGTTCCACGGACGTGTCTACCGGCACCTATACG
Envelope protein
L K G V S Y S L C T A A F T F T K I P A E T L H G T V T V E L Q Y A G T D G P C K V P A Q M A V D M
487 488 489 490 491 492 493 494 495 496 497 498 499 500 501 502 503 504 505 506 507 508 509 510 511 512 513 514 515 516 517 518 519 520 521 522 523 524 525 526 527 528 529 530 531 532 533 534 535 536

5' AGACACTGACCCCAGTGGGCCGGCTGATCACCGCAAATCCCGTGATCACAGAGAGCACCGAGAACTCCAAGATGATGCTGGAGCTGGACCCCCCTTTTGGCGATAGCTACATCGTGATCGGCGTGGGCGAGAAGAAGATCACACACCACT 1800
3' TCTGTGACTGGGGTCACCCGGCCGACTAGTGGCGTTTAGGGCACTAGTGTCTCTCGTGGCTCTTGAGGTTCTACTACGACCTCGACCTGGGGGGAAAACCGCTATCGATGTAGCACTAGCCGCACCCGCTCTTCTTCTAGTGTGTGGTGA
Envelope protein
Q T L T P V G R L I T A N P V I T E S T E N S K M M L E L D P P F G D S Y I V I G V G E K K I T H H
537 538 539 540 541 542 543 544 545 546 547 548 549 550 551 552 553 554 555 556 557 558 559 560 561 562 563 564 565 566 567 568 569 570 571 572 573 574 575 576 577 578 579 580 581 582 583 584 585 586

5' GGCACAGATCTGGCAGCACAATCGGCAAGGCCTTTGAGGCCACCGTGAGGGGAGCAAAGAGGATGGCCGTGCTGGGCGACACCGCATGGGATTTCGGATCCGTGGGAGGCGCCCTGAATTCTCTGGGCAAGGGCATCCACCAGATCTTCG 1950
3' CCGTGTCTAGACCGTCGTGTTAGCCGTTCCGGAAACTCCGGTGGCACTCCCCTCGTTTCTCCTACCGGCACGACCCGCTGTGGCGTACCCTAAAGCCTAGGCACCCTCCGCGGGACTTAAGAGACCCGTTCCCGTAGGTGGTCTAGAAGC
Envelope protein
W H R S G S T I G K A F E A T V R G A K R M A V L G D T A W D F G S V G G A L N S L G K G I H Q I F
587 588 589 590 591 592 593 594 595 596 597 598 599 600 601 602 603 604 605 606 607 608 609 610 611 612 613 614 615 616 617 618 619 620 621 622 623 624 625 626 627 628 629 630 631 632 633 634 635 636

5' GCGCCGCCTTTAAGTCCCTGTTCGGCGGCATGTCTTGGTTTAGCCAGATCCTGATCGGCACACTGCTGATGTGGCTGGGCCTGAACACCAAGAATGGCTCCATCTCTCTGATGTGCCTGGCCCTGGGAGGCGTGCTGATCTTCCTGTCTA 2100
3' CGCGGCGGAAATTCAGGGACAAGCCGCCGTACAGAACCAAATCGGTCTAGGACTAGCCGTGTGACGACTACACCGACCCGGACTTGTGGTTCTTACCGAGGTAGAGAGACTACACGGACCGGGACCCTCCGCACGACTAGAAGGACAGAT
Envelope protein
G A A F K S L F G G M S W F S Q I L I G T L L M W L G L N T K N G S I S L M C L A L G G V L I F L S
637 638 639 640 641 642 643 644 645 646 647 648 649 650 651 652 653 654 655 656 657 658 659 660 661 662 663 664 665 666 667 668 669 670 671 672 673 674 675 676 677 678 679 680 681 682 683 684 685 686

5' CCGCCGTGAGCGCCGACGTGGGATGCTCCGTGGATTTTTCTAAGAAGGAGACACGGTGTGGCACCGGCGTGTTCGTGTACAATGACGTGGAGGCCTGGAGGGATCGCTACAAGTATCACCCTGACAGCCCACGGAGACTGGCAGCAGCCG 2250
3' GGCGGCACTCGCGGCTGCACCCTACGAGGCACCTAAAAAGATTCTTCCTCTGTGCCACACCGTGGCCGCACAAGCACATGTTACTGCACCTCCGGACCTCCCTAGCGATGTTCATAGTGGGACTGTCGGGTGCCTCTGACCGTCGTCGGC
Envelope protein | NS1 protein
T A V S A D V G C S V D F S K K E T R C G T G V F V Y N D V E A W R D R Y K Y H P D S P R R L A A A
687 688 689 690 691 692 693 694 695 696 697 698 699 700 701 702 703 704 705 706 707 708 709 710 711 712 713 714 715 716 717 718 719 720 721 722 723 724 725 726 727 728 729 730 731 732 733 734 735 736

5' TGAAGCAGGCCTGGGAGGATGGCATCTGTGGCATCTCCAGCGTGAGCCGGATGGAGAACATCATGTGGAGAAGCGTGGAGGGCGAGCTGAACGCCATCCTGGAGGAGAATGGAGTGCAGCTGACCGTGGTGGTGGGATCCGTGAAGAATC 2400
3' ACTTCGTCCGGACCCTCCTACCGTAGACACCGTAGAGGTCGCACTCGGCCTACCTCTTGTAGTACACCTCTTCGCACCTCCCGCTCGACTTGCGGTAGGACCTCCTCTTACCTCACGTCGACTGGCACCACCACCCTAGGCACTTCTTAG
NS1 protein
V K Q A W E D G I C G I S S V S R M E N I M W R S V E G E L N A I L E E N G V Q L T V V V G S V K N
737 738 739 740 741 742 743 744 745 746 747 748 749 750 751 752 753 754 755 756 757 758 759 760 761 762 763 764 765 766 767 768 769 770 771 772 773 774 775 776 777 778 779 780 781 782 783 784 785 786

5' CAATGTGGAGGGGACCACAGAGACTGCCAGTGCCTGTGAACGAGCTGCCACACGGATGGAAGGCATGGGGCAAGAGCTATTTTGTGCGGGCCGCCAAGACAAACAATTCCTTCGTGGTGGACGGCGATACCCTGAAGGAGTGCCCCCTGA 2550
3' GTTACACCTCCCCTGGTGTCTCTGACGGTCACGGACACTTGCTCGACGGTGTGCCTACCTTCCGTACCCCGTTCTCGATAAAACACGCCCGGCGGTTCTGTTTGTTAAGGAAGCACCACCTGCCGCTATGGGACTTCCTCACGGGGGACT
NS1 protein
P M W R G P Q R L P V P V N E L P H G W K A W G K S Y F V R A A K T N N S F V V D G D T L K E C P L
787 788 789 790 791 792 793 794 795 796 797 798 799 800 801 802 803 804 805 806 807 808 809 810 811 812 813 814 815 816 817 818 819 820 821 822 823 824 825 826 827 828 829 830 831 832 833 834 835 836

5' AGCACAGAGCCTGGAACAGCTTCCTGGTGGAGGACCACGGCTTCGGCGTGTTTCACACATCCGTGTGGCTGAAGGTGAGAGAGGACTACTCTCTGGAGTGTGATCCAGCCGTGATCGGAACCGCAGTGAAGGGCAAGGAGGCAGTGCACA 2700
3' TCGTGTCTCGGACCTTGTCGAAGGACCACCTCCTGGTGCCGAAGCCGCACAAAGTGTGTAGGCACACCGACTTCCACTCTCTCCTGATGAGAGACCTCACACTAGGTCGGCACTAGCCTTGGCGTCACTTCCCGTTCCTCCGTCACGTGT
NS1 protein
K H R A W N S F L V E D H G F G V F H T S V W L K V R E D Y S L E C D P A V I G T A V K G K E A V H
837 838 839 840 841 842 843 844 845 846 847 848 849 850 851 852 853 854 855 856 857 858 859 860 861 862 863 864 865 866 867 868 869 870 871 872 873 874 875 876 877 878 879 880 881 882 883 884 885 886

5' GCGACCTGGGCTATTGGATCGAGTCCGAGAAGAATGATACATGGCGGCTGAAGAGAGCCCACCTGATCGAGATGAAGACCTGCGAGTGGCCCAAGTCTCACACACTGTGGACCGACGGCATCGAGGAGAGCGATCTGATCATCCCTAAGT 2850
3' CGCTGGACCCGATAACCTAGCTCAGGCTCTTCTTACTATGTACCGCCGACTTCTCTCGGGTGGACTAGCTCTACTTCTGGACGCTCACCGGGTTCAGAGTGTGTGACACCTGGCTGCCGTAGCTCCTCTCGCTAGACTAGTAGGGATTCA
NS1 protein
S D L G Y W I E S E K N D T W R L K R A H L I E M K T C E W P K S H T L W T D G I E E S D L I I P K
887 888 889 890 891 892 893 894 895 896 897 898 899 900 901 902 903 904 905 906 907 908 909 910 911 912 913 914 915 916 917 918 919 920 921 922 923 924 925 926 927 928 929 930 931 932 933 934 935 936

5' CTCTGGCCGGCCCACTGAGCCACCACAACACAAGGGAGGGCTACCGCACCCAGATGAAGGGCCCTTGGCACAGCGAGGAGCTGGAGATCAGGTTTGAGGAGTGCCCAGGCACCAAGGTGCACGTGGAGGAGACATGTGGAACCCGGGGAC 3000
3' GAGACCGGCCGGGTGACTCGGTGGTGTTGTGTTCCCTCCCGATGGCGTGGGTCTACTTCCCGGGAACCGTGTCGCTCCTCGACCTCTAGTCCAAACTCCTCACGGGTCCGTGGTTCCACGTGCACCTCCTCTGTACACCTTGGGCCCCTG
NS1 protein
S L A G P L S H H N T R E G Y R T Q M K G P W H S E E L E I R F E E C P G T K V H V E E T C G T R G
937 938 939 940 941 942 943 944 945 946 947 948 949 950 951 952 953 954 955 956 957 958 959 960 961 962 963 964 965 966 967 968 969 970 971 972 973 974 975 976 977 978 979 980 981 982 983 984 985 986

5' CATCCCTGAGATCTACAACCGCCTCTGGCAGAGTGATCGAGGAGTGGTGCTGTCGCGAGTGCACCATGCCACCACTGAGCTTCCGCGCAAAGGACGGCTGTTGGTATGGCATGGAGATTCGCCCACGCAAGGAACCTGAAAGCAACCTGG 3150
3' GTAGGGACTCTAGATGTTGGCGGAGACCGTCTCACTAGCTCCTCACCACGACAGCGCTCACGTGGTACGGTGGTGACTCGAAGGCGCGTTTCCTGCCGACAACCATACCGTACCTCTAAGCGGGTGCGTTCCTTGGACTTTCGTTGGACC
NS1 protein
P S L R S T T A S G R V I E E W C C R E C T M P P L S F R A K D G C W Y G M E I R P R K E P E S N L
987 988 989 990 991 992 993 994 995 996 997 998 999 1000 1001 1002 1003 1004 1005 1006 1007 1008 1009 1010 1011 1012 1013 1014 1015 1016 1017 1018 1019 1020 1021 1022 1023 1024 1025 1026 1027 1028 1029 1030 1031 1032 1033 1034 1035 1036

                              AatII
5' TCCGCTCTATGGTGACAGCATAGCCGACGTCGCGATCGCACTAGTGTGAAATAGACATCAG 3211
3' AGGCGAGATACCACTGTCGTATCGGCTGCAGCGCTAGCGTGATCACACTTTATCTGTAGTC
NS1 protein
V R S M V T A .
1037 1038 1039 1040 1041 1042 1043 1044
```

MluI

5' CAGAGTGATACGCGTGGCCACCATGGAGACAGACACACTCCTGCTATGGGTACTGCTGCTCTGGGTTCCAGGTTCCACTGGTGACGTGGGATGCTCCGTGGATTTTTCTAAGAAGGAGACACGGTGTGGCACCGGCGTGTTCGTGTACAA 150

3' GTCTCACTATGCGCACCGGTGGTACCTCTGTCTGTGTGAGGACGATACCCATGACGACGAGACCCAAGGTCCAAGGTGACCACTGCACCCTACGAGGCACCTAAAAAGATTCTTCCTCTGTGCCACACCGTGGCCGCACAAGCACATGTT

NS1 protein

Ig Kappa Signal peptide

M E T D T L L L W V L L L W V P G S T G D V G C S V D F S K K E T R C G T G V F V Y N

1 2 3 4 5 6 7 8 9 10 11 12 13 14 15 16 17 18 19 20 21 22 23 24 25 26 27 28 29 30 31 32 33 34 35 36 37 38 39 40 41 42 43

5' TGACGTGGAGGCCTGGAGGGATCGCTACAAGTATCACCCTGACAGCCCACGGAGACTGGCAGCAGCCGTGAAGCAGGCCTGGGAGGATGGCATCTGTGGCATCTCCAGCGTGAGCCGGATGGAGAACATCATGTGGAGAAGCGTGGAGGG 300

3' ACTGCACCTCCGGACCTCCCTAGCGATGTTCATAGTGGGACTGTCGGGTGCCTCTGACCGTCGTCGGCACTTCGTCCGGACCCTCCTACCGTAGACACCGTAGAGGTCGCACTCGGCCTACCTCTTGTAGTACACCTCTTCGCACCTCCC

NS1 protein

D V E A W R D R Y K Y H P D S P R R L A A A V K Q A W E D G I C G I S S V S R M E N I M W R S V E G

44 45 46 47 48 49 50 51 52 53 54 55 56 57 58 59 60 61 62 63 64 65 66 67 68 69 70 71 72 73 74 75 76 77 78 79 80 81 82 83 84 85 86 87 88 89 90 91 92 93

5' CGAGCTGAACGCCATCCTGGAGGAGAATGGAGTGCAGCTGACCGTGGTGGTGGGATCCGTGAAGAATCCAATGTGGAGGGGACCACAGAGACTGCCAGTGCCTGTGAACGAGCTGCCACACGGATGGAAGGCATGGGGCAAGAGCTATTT 450

3' GCTCGACTTGCGGTAGGACCTCCTCTTACCTCACGTCGACTGGCACCACCACCCTAGGCACTTCTTAGGTTACACCTCCCCTGGTGTCTCTGACGGTCACGGACACTTGCTCGACGGTGTGCCTACCTTCCGTACCCCGTTCTCGATAAA

NS1 protein

E L N A I L E E N G V Q L T V V V G S V K N P M W R G P Q R L P V P V N E L P H G W K A W G K S Y F

94 95 96 97 98 99 100 101 102 103 104 105 106 107 108 109 110 111 112 113 114 115 116 117 118 119 120 121 122 123 124 125 126 127 128 129 130 131 132 133 134 135 136 137 138 139 140 141 142 143

ArsI'

5' TGTGCGGGCCGCCAAGACAAACAATTCCTTCGTGGTGGACGGCGATACCCTGAAGGAGTGCCCCCTGAAGCACAGAGCCTGGAACAGCTTCCTGGTGGAGGACCACGGCTTCGGCGTGTTTCACACATCCGTGTGGCTGAAGGTGAGAGA 600

3' ACACGCCCGGCGGTTCTGTTTGTTAAGGAAGCACCACCTGCCGCTATGGGACTTCCTCACGGGGGACTTCGTGTCTCGGACCTTGTCGAAGGACCACCTCCTGGTGCCGAAGCCGCACAAAGTGTGTAGGCACACCGACTTCCACTCTCT

NS1 protein

V R A A K T N N S F V V D G D T L K E C P L K H R A W N S F L V E D H G F G V F H T S V W L K V R E

144 145 146 147 148 149 150 151 152 153 154 155 156 157 158 159 160 161 162 163 164 165 166 167 168 169 170 171 172 173 174 175 176 177 178 179 180 181 182 183 184 185 186 187 188 189 190 191 192 193

5' GGACTACTCTCTGGAGTGTGATCCAGCCGTGATCGGAACCGCAGTGAAGGGCAAGGAGGCAGTGCACAGCGACCTGGGCTATTGGATCGAGTCCGAGAAGAATGATACATGGCGGCTGAAGAGAGCCCACCTGATCGAGATGAAGACCTG 750

3' CCTGATGAGAGACCTCACACTAGGTCGGCACTAGCCTTGGCGTCACTTCCCGTTCCTCCGTCACGTGTCGCTGGACCCGATAACCTAGCTCAGGCTCTTCTTACTATGTACCGCCGACTTCTCTCGGGTGGACTAGCTCTACTTCTGGAC

NS1 protein

D Y S L E C D P A V I G T A V K G K E A V H S D L G Y W I E S E K N D T W R L K R A H L I E M K T C

194 195 196 197 198 199 200 201 202 203 204 205 206 207 208 209 210 211 212 213 214 215 216 217 218 219 220 221 222 223 224 225 226 227 228 229 230 231 232 233 234 235 236 237 238 239 240 241 242 243

5' CGAGTGGCCCAAGTCTCACACACTGTGGACCGACGGCATCGAGGAGAGCGATCTGATCATCCCTAAGTCTCTGGCCGGCCCACTGAGCCACCACAACACAAGGGAGGGCTACCGCACCCAGATGAAGGGCCCTTGGCACAGCGAGGAGCT 900

3' GCTCACCGGGTTCAGAGTGTGTGACACCTGGCTGCCGTAGCTCCTCTCGCTAGACTAGTAGGGATTCAGAGACCGGCCGGGTGACTCGGTGGTGTTGTGTTCCCTCCCGATGGCGTGGGTCTACTTCCCGGGAACCGTGTCGCTCCTCGA

NS1 protein

E W P K S H T L W T D G I E E S D L I I P K S L A G P L S H H N T R E G Y R T Q M K G P W H S E E L

244 245 246 247 248 249 250 251 252 253 254 255 256 257 258 259 260 261 262 263 264 265 266 267 268 269 270 271 272 273 274 275 276 277 278 279 280 281 282 283 284 285 286 287 288 289 290 291 292 293

5' GGAGATCAGGTTTGAGGAGTGCCCAGGCACCAAGGTGCACGTGGAGGAGACATGTGGAACCCGGGGACCATCCCTGAGATCTACAACCGCCTCTGGCAGAGTGATCGAGGAGTGGTGCTGTCGCGAGTGCACCATGCCACCACTGAGCTT 1050

3' CCTCTAGTCCAAACTCCTCACGGGTCCGTGGTTCCACGTGCACCTCCTCTGTACACCTTGGGCCCCTGGTAGGGACTCTAGATGTTGGCGGAGACCGTCTCACTAGCTCCTCACCACGACAGCGCTCACGTGGTACGGTGGTGACTCGAA

NS1 protein

E I R F E E C P G T K V H V E E T C G T R G P S L R S T T A S G R V I E E W C C R E C T M P P L S F

294 295 296 297 298 299 300 301 302 303 304 305 306 307 308 309 310 311 312 313 314 315 316 317 318 319 320 321 322 323 324 325 326 327 328 329 330 331 332 333 334 335 336 337 338 339 340 341 342 343

AatII AsiSI

5' CCGCGCAAAGGACGGCTGTTGGTATGGCATGGAGATTCGCCCACGCAAGGAACCTGAAAGCAACCTGGTCCGCTCTATGGTGACAGCATAGCCGACGTCTTGCGATCGCGTGCGAGAGGCCAGAACAACATCCGCCTACC 1190

3' GGCGCGTTTCCTGCCGACAACCATACCGTACCTCTAAGCGGGTGCGTTCCTTGGACTTTCGTTGGACCAGGCGAGATACCACTGTCGTATCGGCTGCAGAACGCTAGCGCACGCTCTCCGGTCTTGTTGTAGGCGGATGG

NS1 protein

R A K D G C W Y G M E I R P R K E P E S N L V R S M V T A .

344 345 346 347 348 349 350 351 352 353 354 355 356 357 358 359 360 361 362 363 364 365 366 367 368 369 370 371 372 373

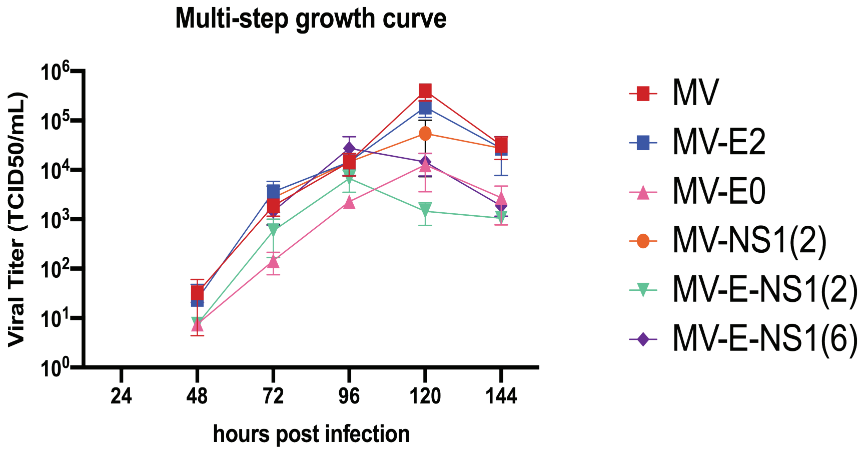
Multi-step growth curve
Viral Titer (TCID50/mL)
10^0
10^1
10^2
10^3
10^4
10^5
10^6
24
48
72
96
120
144
hours post infection
MV
MV-E2
MV-E0
MV-NS1(2)
MV-E-NS1(2)
MV-E-NS1(6)

A

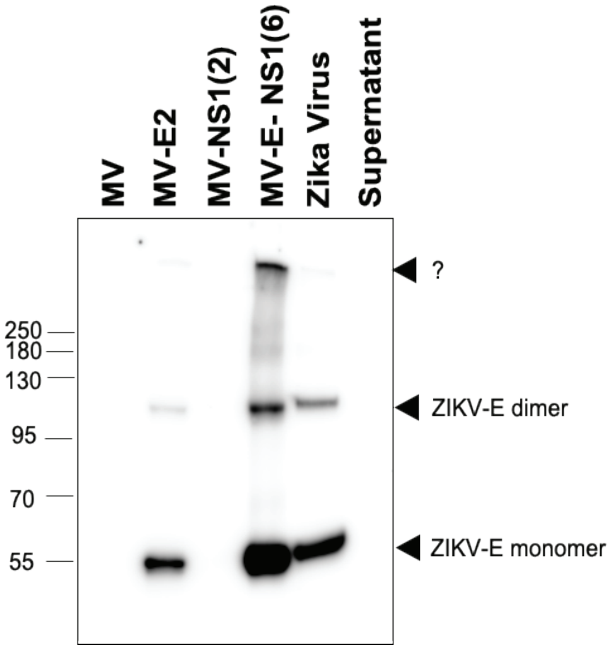
MV
MV-E2
MV-NS1(2)
MV-E- NS1(6)
Zika Virus
Supernatant
250
180
130
95
70
55
?
ZIKV-E dimer
ZIKV-E monomer

B

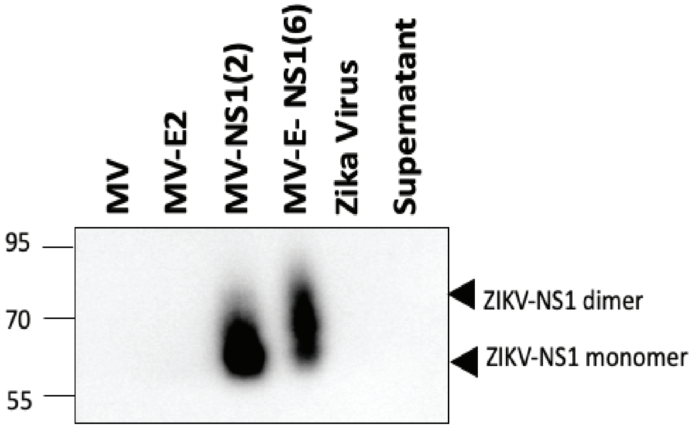
MV
MV-E2
MV-NS1(2)
MV-E- NS1(6)
Zika Virus
Supernatant
95
70
55
ZIKV-NS1 dimer
ZIKV-NS1 monomer

```
BglII
5' TCGTGGTGCCAGATCTCACAGAGCCGCCATCTATACCGCCGAGATCCACAAGTCTCTGAGCACCAACCTGGACGTGACAAATTCCATCGAGCACCAGGTGAAGGATGTGC 110
3' AGCACCACGGTCTAGAGTGTCTCGGCGGTAGATATGGCGGCTCTAGGTGTTCAGAGACTCGTGGTTGGACCTGCACTGTTTAAGGTAGCTCGTGGTCCACTTCCTACACG
co MV Hemagglutinin
H R A A I Y T A E I H K S L S T N L D V T N S I E H Q V K D V
1 2 3 4 5 6 7 8 9 10 11 12 13 14 15 16 17 18 19 20 21 22 23 24 25 26 27 28 29 30 31

5' TGACACCACTGTTTAAGATCATCGGCGACGAGGTCGGCCTGAGGACCCCACAGCGGTTCACCGACCTGGTGAAGCTGATCTCTGATAAGATCAAGTTTCTGAACCCCGAT 220
3' ACTGTGGTGACAAATTCTAGTAGCCGCTGCTCCAGCCGGACTCCTGGGGTGTCGCCAAGTGGCTGGACCACTTCGACTAGAGACTATTCTAGTTCAAAGACTTGGGGCTA
co MV Hemagglutinin
L T P L F K I I G D E V G L R T P Q R F T D L V K L I S D K I K F L N P D
32 33 34 35 36 37 38 39 40 41 42 43 44 45 46 47 48 49 50 51 52 53 54 55 56 57 58 59 60 61 62 63 64 65 66 67 68

5' CGGGAGTACGACTTCCGGGACCTGACCTGGTGCATCAATCCCCCTGAGCGGATCAAGCTGGACTACGATCAGTATTGTGCCGACGTGGCCGCCGAGGAGCTGATGAACGC 330
3' GCCCTCATGCTGAAGGCCCTGGACTGGACCACGTAGTTAGGGGGACTCGCCTAGTTCGACCTGATGCTAGTCATAACACGGCTGCACCGGCGGCTCCTCGACTACTTGCG
co MV Hemagglutinin
R E Y D F R D L T W C I N P P E R I K L D Y D Q Y C A D V A A E E L M N A
69 70 71 72 73 74 75 76 77 78 79 80 81 82 83 84 85 86 87 88 89 90 91 92 93 94 95 96 97 98 99 100 101 102 103 104 105

5' CCTGGTGAATAGCACCCTGCTGGAGACAAGAACCACAAACCAGTTTCTGGCCGTGTCCAAGGGCAATTGCTCTGGCCCTACCACAATCAGGGGCCAGTTCTCCAACATGT 440
3' GGACCACTTATCGTGGGACGACCTCTGTTCTTGGTGTTTGGTCAAAGACCGGCACAGGTTCCCGTTAACGAGACCGGGATGGTGTTAGTCCCCGGTCAAGAGGTTGTACA
co MV Hemagglutinin
L V N S T L L E T R T T N Q F L A V S K G N C S G P T T I R G Q F S N M
106 107 108 109 110 111 112 113 114 115 116 117 118 119 120 121 122 123 124 125 126 127 128 129 130 131 132 133 134 135 136 137 138 139 140 141

5' CCCTGTCTCTGCTGGATCTGTACCTGGGCCGCGGCTATAATGTGAGCTCCATCGTGACCATGACATCTCAGGGCATGTACGGCGGCACCTATCTGGTGGAGAAGCCAAAC 550
3' GGGACAGAGACGACCTAGACATGGACCCGGCGCCGATATTACACTCGAGGTAGCACTGGTACTGTAGAGTCCCGTACATGCCGCCGTGGATAGACCACCTCTTCGGTTTG
co MV Hemagglutinin
S L S L L D L Y L G R G Y N V S S I V T M T S Q G M Y G G T Y L V E K P N
142 143 144 145 146 147 148 149 150 151 152 153 154 155 156 157 158 159 160 161 162 163 164 165 166 167 168 169 170 171 172 173 174 175 176 177 178

5' CTGTCTAGCAAGCGGAGCGAGCTGTCCCAGCTGTCTATGTACAGGGTGTTTGAAGTGGGCGTGATCAGGAACCCTGGCCTGGGAGCCCCAGTGTTCCACATGACCAATTA 660
3' GACAGATCGTTCGCCTCGCTCGACAGGGTCGACAGATACATGTCCCACAAACTTCACCCGCACTAGTCCTTGGGACCGGACCCTCGGGGTCACAAGGTGTACTGGTTAAT
co MV Hemagglutinin
L S S K R S E L S Q L S M Y R V F E V G V I R N P G L G A P V F H M T N Y
179 180 181 182 183 184 185 186 187 188 189 190 191 192 193 194 195 196 197 198 199 200 201 202 203 204 205 206 207 208 209 210 211 212 213 214 215

5' TCTGGAGCAGCCAGTGAGCAACGATCTGTCCAATTGCATGGTGGCCCTGGGAGAGCTGAAGCTGGCCGCCCTGTGCCACGGCGAGGACTCTATCACCATCCCATACCAGG 770
3' AGACCTCGTCGGTCACTCGTTGCTAGACAGGTTAACGTACCACCGGGACCCTCTCGACTTCGACCGGCGGGACACGGTGCCGCTCCTGAGATAGTGGTAGGGTATGGTCC
co MV Hemagglutinin
L E Q P V S N D L S N C M V A L G E L K L A A L C H G E D S I T I P Y Q
216 217 218 219 220 221 222 223 224 225 226 227 228 229 230 231 232 233 234 235 236 237 238 239 240 241 242 243 244 245 246 247 248 249 250 251

5' GAAGCGGCAAGGGCGTGTCCTTTCAGCTGGTGAAGCTGGGCGTGTGGAAGTCCCCTACCGACATGCAGTCTTGGGTGCCCCTGAGCACAGACGATCCTGTGATCGATCGG 880
3' CTTCGCCGTTCCCGCACAGGAAAGTCGACCACTTCGACCCGCACACCTTCAGGGGATGGCTGTACGTCAGAACCCACGGGGACTCGTGTCTGCTAGGACACTAGCTAGCC
co MV Hemagglutinin
G S G K G V S F Q L V K L G V W K S P T D M Q S W V P L S T D D P V I D R
252 253 254 255 256 257 258 259 260 261 262 263 264 265 266 267 268 269 270 271 272 273 274 275 276 277 278 279 280 281 282 283 284 285 286 287 288

5' CTGTATCTGTCCTCTCACAGAGGCGTGATCGCCGACAATCAGGCCAAGTGGGCCGTGCCAACCACAAGGACCGACGATAAGCTGCGCATGGAGACATGCTTTCAGCAGGC 990
3' GACATAGACAGGAGAGTGTCTCCGCACTAGCGGCTGTTAGTCCGGTTCACCCGGCACGGTTGGTGTTCCTGGCTGCTATTCGACGCGTACCTCTGTACGAAAGTCGTCCG
co MV Hemagglutinin
L Y L S S H R G V I A D N Q A K W A V P T T R T D D K L R M E T C F Q Q A
289 290 291 292 293 294 295 296 297 298 299 300 301 302 303 304 305 306 307 308 309 310 311 312 313 314 315 316 317 318 319 320 321 322 323 324 325

5' CTGTAAGGGCAAGATCCAGGCCCTGTGCGAGAACCCTGAGTGGGCACCACTGAAGGATAATCGGATCCCAAGCTACGGCGTGCTGTCTGTGGACCTGAGCCTGACCGTGG 1100
3' GACATTCCCGTTCTAGGTCCGGGACACGCTCTTGGGACTCACCCGTGGTGACTTCCTATTAGCCTAGGGTTCGATGCCGCACGACAGACACCTGGACTCGGACTGGCACC
co MV Hemagglutinin
C K G K I Q A L C E N P E W A P L K D N R I P S Y G V L S V D L S L T V
326 327 328 329 330 331 332 333 334 335 336 337 338 339 340 341 342 343 344 345 346 347 348 349 350 351 352 353 354 355 356 357 358 359 360 361

5' AGCTGAAGATCAAGATCGCCTCCGGCTTCGGCCCCCTGATCACACACGGCTCCGGCATGGATCTGTACAAGTCTAACCACAACAACGTGTACTGGCTGACCATCCCACCC 1210
3' TCGACTTCTAGTTCTAGCGGAGGCCGAAGCCGGGGGACTAGTGTGTGCCGAGGCCGTACCTAGACATGTTCAGATTGGTGTTGTTGCACATGACCGACTGGTAGGGTGGG
co MV Hemagglutinin
E L K I K I A S G F G P L I T H G S G M D L Y K S N H N N V Y W L T I P P
362 363 364 365 366 367 368 369 370 371 372 373 374 375 376 377 378 379 380 381 382 383 384 385 386 387 388 389 390 391 392 393 394 395 396 397 398

5' ATGAAGAACCTGGCCCTGGGCGTGATCAATACACTGGAGTGGATCCCACGGTTCAAGGTGAGCCCCTACCTGTTCACCGTGCCTATCAAGGAGGCAGGAGAGGACTGCCA 1320
3' TACTTCTTGGACCGGGACCCGCACTAGTTATGTGACCTCACCTAGGGTGCCAAGTTCCACTCGGGGATGGACAAGTGGCACGGATAGTTCCTCCGTCCTCTCCTGACGGT
co MV Hemagglutinin
M K N L A L G V I N T L E W I P R F K V S P Y L F T V P I K E A G E D C H
399 400 401 402 403 404 405 406 407 408 409 410 411 412 413 414 415 416 417 418 419 420 421 422 423 424 425 426 427 428 429 430 431 432 433 434 435

5' CGCACCAACATATCTGCCTGCCGAGGTGGACGGCGATGTGAAGCTGAGCTCCAACCTGGTCATCCTGCCAGGACAGGACCTGCAGTACGTGCTGGCCACCTATGATACAA 1430
3' GCGTGGTTGTATAGACGGACGGCTCCACCTGCCGCTACACTTCGACTCGAGGTTGGACCAGTAGGACGGTCCTGTCCTGGACGTCATGCACGACCGGTGGATACTATGTT
co MV Hemagglutinin
A P T Y L P A E V D G D V K L S S N L V I L P G Q D L Q Y V L A T Y D T
436 437 438 439 440 441 442 443 444 445 446 447 448 449 450 451 452 453 454 455 456 457 458 459 460 461 462 463 464 465 466 467 468 469 470 471

5' GCAGGGTGGAGCACGCCGTGGTGTACTACGTGTACAGCCCTTCCCGGTCTTTCAGCTACTTTTATCCATTCAGACTGCCCATCAAGGGCGTGCCAATCGAGCTGCAGGTG 1540
3' CGTCCCACCTCGTGCGGCACCACATGATGCACATGTCGGGAAGGGCCAGAAAGTCGATGAAAATAGGTAAGTCTGACGGGTAGTTCCCGCACGGTTAGCTCGACGTCCAC
co MV Hemagglutinin
S R V E H A V V Y Y V Y S P S R S F S Y F Y P F R L P I K G V P I E L Q V
472 473 474 475 476 477 478 479 480 481 482 483 484 485 486 487 488 489 490 491 492 493 494 495 496 497 498 499 500 501 502 503 504 505 506 507 508

5' GAGTGTTTTACCTGGGATCAGAAGCTGTGGTGCCGCCACTTCTGCGTGCTGGCCGACTCCGAGTCTGGAGGACACATCACACACTCCGGCATGGTCGGGATGGGGGTGTC 1650
3' CTCACAAAATGGACCCTAGTCTTCGACACCACGGCGGTGAAGACGCACGACCGGCTGAGGCTCAGACCTCCTGTGTAGTGTGTGAGGCCGTACCAGCCCTACCCCCACAG
co MV Hemagglutinin
E C F T W D Q K L W C R H F C V L A D S E S G G H I T H S G M V G M G V S
509 510 511 512 513 514 515 516 517 518 519 520 521 522 523 524 525 526 527 528 529 530 531 532 533 534 535 536 537 538 539 540 541 542 543 544 545

NotI
5' TTGCACTGTCACTCGGGAGGACGGCACAAATAGAAGGTAGgcggccgccgctcgaga 1707
3' AACGTGACAGTGAGCCCTCCTGCCGTGTTTATCTTCCATCcgccggcggcgagctct
co MV Hemagglutinin
C T V T R E D G T N R R .
546 547 548 549 550 551 552 553 554 555 556 557 558
```
